# Transient hypoxia followed by progressive reoxygenation is required for efficient skeletal muscle repair through Rev-ERBα modulation

**DOI:** 10.1101/2024.05.02.592180

**Authors:** Marie Quétin, Audrey Der Vartanian, Christelle Dubois, Juliette Berthier, Marine Ledoux, Stéphanie Michineau, Bernadette Drayton-Libotte, Athanassia Sotiropoulos, Frédéric Relaix, Marianne Gervais

## Abstract

Muscle stem cells (MuSCs) are essential for skeletal muscle repair. Following injury, MuSCs reside in low oxygen environments until muscle fibers and vascularization are restablished. The dynamics of oxygen levels during the regenerative process and its impact on muscle repair has been underappreciated. We confirm that muscle repair is initiated in a low oxygen environment followed by gradual reoxygenation. Strikingly, when muscle reoxygenation is limited by keeping mice under systemic hypoxia, muscle repair is impaired and leads to the formation of hypotrophic myofibers. *In vivo*, sustained hypoxia decreases the ability of MuSCs to differentiate and fuse independently of HIF-1α. Prolonged hypoxia specifically affects the circadian clock by increasing *Rev-erbα* expression in MuSCs. Using pharmacological tools, we demonstrate that Rev-ERBα negatively regulates myogenesis by reducing late myogenic cell fusion under prolonged hypoxia. Our results underscore the critical role of progressive muscle reoxygenation after transient hypoxia in coordinating proper myogenesis through Rev-ERBα.

## INTRODUCTION

Skeletal muscle exhibits robust growth^1^ and regenerative capacity^2^ which depend on the muscle stem cells (MuSCs), also known as muscle satellite cells. In adult muscle, MuSCs are maintained into quiescence and express the PAX7 transcription factor^3^. Upon muscle damage, MuSCs activate and proliferate co-expressing PAX7 and MYOD before differentiating by downregulating PAX7 and inducing MYOGENIN (also known as MYOG) to generate new myofibers^4^. Indeed, differentiated myoblasts may either fuse to each other forming *de novo* myofibers or fuse with existing myotubes in the injured muscle. This additional fusion of myoblasts, called myonuclear accretion, ensures myofiber growth by increasing myonuclei number^5^. The accomplishment of this fusion process requires cell-fusion molecules, especially MYOMAKER, which is expressed in both fusogenic cells^6^. In parallel, a subset population of activated MuSCs downregulate MYOD and exit the cell cycle to self-renew the pool of quiescent PAX7^+^ MuSCs for future needs^7^.

Skeletal myogenesis and muscle vascularization are coupled during postnatal muscle growth^8^ and muscle repair^9^. Vascular alterations result in reduced oxygen (O_2_) levels, disrupting cell homeostasis and contributing to many diseases, especially through Hypoxia Inducible Factors (HIFs) and its most studied isoform HIF-1α^10^. In skeletal muscles, HIF-1α level increases during exercise^11^, after systemic hypoxia^12^ or during muscle injury in rodents^13^. Although the role of hypoxia, HIFs and HIF-inducible genes are well established in angiogenesis^14^, their impact on MuSC specification and myogenesis remains controversial. *In vitro*, most studies agree that hypoxia promotes MuSC quiescence thought HIF-1α and Notch signaling^15,16,17^, while the impact of hypoxia and HIFs on myogenic cell proliferation and differentiation remains controversial^18,19,20^. *In vivo*, Majmundar and colleagues show that HIF-1α in MuSCs negatively regulates myogenesis by decreasing myogenic differentiation^21^.

Molecular adaptations of skeletal muscle and satellite cells to hypoxia may also be HIF-independent. Many evidences suggest that hypoxia decreases MyoD/MyoG expressions and subsequently myogenic cell differentiation though the induction of myostatin^22^, Bhlhe40^23^ and HDAC9^20^. Hypoxia can also alter myogenic differentiation and myotube formation by inhibiting p21 (as known as p21 and CDKN1A) that leads to an accumulation of the retinoblastoma protein Rb^24^. Moreover, it has been demonstrated that the repression of PI3K-Akt-mTOR pathway at low O_2_ levels decreases the differentiation capacity of MuSCs^25^ and may leads to an impairment of myofiber formation and growth^18^.

These studies reveal a wide range of factors and signaling pathways regulating myogenesis under hypoxia. However, it remains unclear how oxygen level variation *in vivo* influences myogenic cell fate for effective muscle repair.

In this study, we established that skeletal muscle regeneration after cardiotoxin-induced injury is initiated in a low oxygen environment and is followed by a progressive reoxygenation. To investigate whether muscle reoxygenation following transient hypoxia is crucial for effective muscle repair, we blunted muscle reoxygenation by maintaining mice under systemic hypoxia. Our results demonstrate that prolonged hypoxia impairs regenerative myogenesis and leads to long-term hypotrophic regenerating phenotype, by reducing the differentiation and fusion capacities of MuSCs. On isolated myofibers, we show that prolonged hypoxia (1%O_2_) promotes the self-renewal but limits the differentiation of MuSCs, as compared to physioxia (8%O_2_). In contrast, transient hypoxia followed by progressive reoxygenation (1 to 8%O_2_) enhances MuSC self-renewal and restores their differentiation capacity. Using *Pax7^CreERT^*^2^*^/+^*;*HIF1α^flox/flox^* mutant mice, we established that renewal of the MuSC pool following transient hypoxia and progressive reoxygenation is HIF-1α-dependent. However, their differentiation and fusion potentials are independent of HIF-1α. Transcriptomic analysis on FACS-sorted MuSCs from muscles underdoing hypoxia *versus* normoxia during regeneration, confirmed by *in vitro* experiments, revealed that the upregulation of Rev-Erbα under prolonged hypoxia controls the late myogenic program of MuSCs.

## RESULTS

### Coordinated myo-angiogenesis during muscle repair is initiated in a hypoxic environment followed by progressive reoxygenation

We analyzed the spatio-temporal patterns of myogenesis, vascularization muscle oxygenation and expression of HIF target genes during skeletal muscle repair in mice that received an intramuscular cardiotoxin (CTX) injection in the *Tibialis Anterior* (TA) muscle (*Fig. 1A*). CTX-induced muscle injury led to disorganized skeletal muscle architecture (*Fig. 1B*) characterized by a drastic drop in capillaries density and fiber size at 5 days post-injury (dpi*; Fig. 1B and 1C*). This phenomenon is associated with a concomitant mobilization of PAX7+ cells to ensure muscle repair (*Fig. 1D*), as already described^26^. From 5 to 28 dpi, the increase in myofiber size correlated with muscle revascularization, demonstrating synchronous myo-angiogenic coupling during muscle repair (*Fig. 1B and 1C*). Restoration of the microvascular network during muscle repair inversely correlates with PAX7+ cell number and their progressive return into quiescence (*Fig. 1B and 1D*). To assess whether alterations of the vascular network following muscle injury are associated with reduced oxygen levels *in situ*, we evaluated the kinetics of partial oxygen tension during muscle repair using a hypoxia probe, named pimonidazole (*Fig. 1A*). When injected *in vivo*, this probe is reductively activated in hypoxic cells where it forms covalent protein adducts that are revealed by immunofluorescence^26^. Strikingly, the most abundant and intense pimonidazole staining is detected on CTX-injured TAs at 5 dpi, indicating that myogenic cell expansion is initiated in a hypoxic environment *in situ* (*Fig. 1D-1F*). From 5 to 28 dpi, pimonidazole adduct intensity gradually declined, demonstrating a progressive reoxygenation after transient hypoxia during muscle repair (*Fig. 1E and 1F*) that correlates with progressive restoration of the vascular network (*Fig. 1C*) and MuSC return into quiescence (*Fig. 1B and 1D*). To confirm the hypoxic status of muscle cells *in vivo,* we monitored the early activation of the HIF signaling pathway. We observed increased expression of HIF-inducible target genes, *Lysyl oxidase homolog 2* (*Loxl-2*), *platelet derived growth factor subunit B* (*Pdgfb*) and *Angiopoietin-2* (*Ang2*) the first week following muscle injury (5-7 dpi), with a return to normal at 14 dpi (*Fig. 1G*).

**Figure 1:**
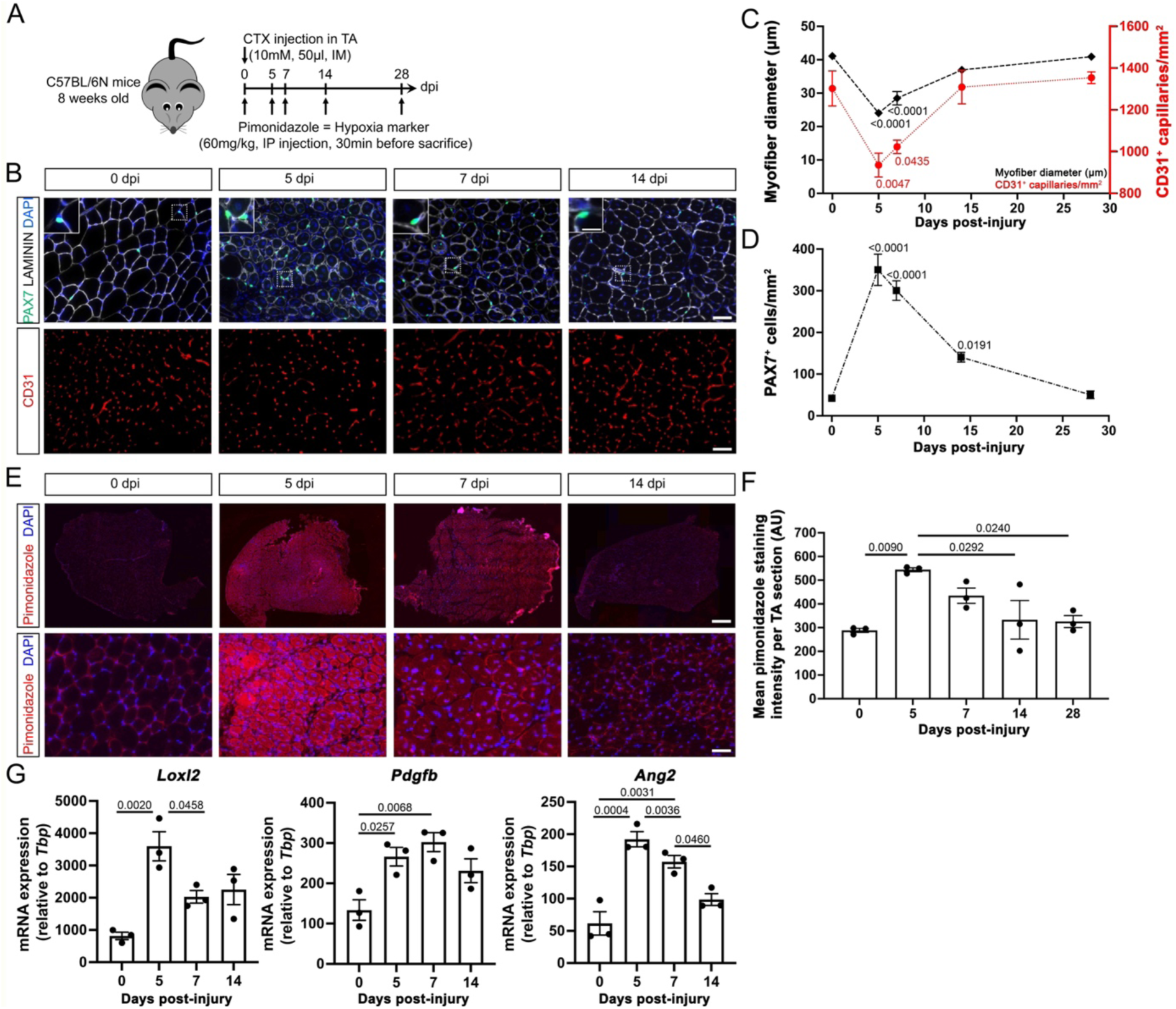
Coordinated myo-angiogenesis during muscle repair is initiated in a hypoxic environment followed by progressive reoxygenation. **A.** Experimental design. Acute muscle injury was induced by cardiotoxin (CTX) intramuscular (IM) injection in the tibialis anterior (TA) of 8-week-old wild-type mice. Intraperitoneal (IP) injection of pimonidazole hypoxic probe were performed during the time course of muscle regeneration at 0, 5, 7, 14 and 28 days post-injury (dpi). **B.** Representative pictures of PAX7 (green), LAMININ (white), CD31 (red) and nuclei (DAPI, blue) staining on CTX-injured TA cross-sections. Scale bars: 50μm (overviews) and 12.5μm (insets). **C.** Quantification of myofiber diameter (μm) and capillaries density (CD31^+^cells/mm^2^) on CTX-injured TA cross-sections. **D.** Quantification of PAX7^+^ cells/mm^2^ on CTX-injured TAs cross-sections. **E.** Representative pictures of pimonidazole staining (red) on CTX-injured TA cross-sections. Scale bars: 200μm (upper panel) and 50μm (lower panel). **F.** Quantification of pimonidazole staining intensity (arbitrary unit, AU) on CTX-injured TA cross-sections. **G.** Histograms showing the relative expression of HIF/HRE target genes, *Ang2 (*angiopoietin 2), *Loxl2 (*lysyl oxidase like 2) and *Pdgfb* (platelet-derived growth factor subunit B) normalized to *Tbp* (TATA-box Binding Protein) on CTX-injured TA extracts. **<u>Statistics</u>:** Results are expressed as means ± SEM. For **C** and **D.** One-way ANOVA followed by Dunnet’s post-test versus 0 dpi. n=3-4 biologically independent samples per time point. For **F** and **G.** One-way ANOVA followed by Tukey’s post-test. n=3 biologically independent samples per time point.

We therefore conclude that *in vivo* post-injury muscle repair is associated with myo-angiogenic coupling that is initiated in a low oxygen environment followed by progressive muscle reoxygenation.

### Blocking muscle reoxygenation impairs skeletal muscle repair and leads to a hypotrophic myofiber phenotype

To address whether progressive reoxygenation after transient hypoxia is required for muscle repair *in vivo*, we tested the impact of systemic hypoxia exposure during skeletal muscle regeneration. TA muscles of adult mice were injured with CTX, and mice were immediately housed in a normal (21%O_2_) or hypoxic (10%O_2_ inhaled) normobaric atmosphere for 28 days (*Fig. 2A*). *In vivo* systemic hypoxia was validated in mice exposed to low oxygen concentration by the significant increase of their hematocrit level from 7dpi to 28 dpi and capillary density at 28 dpi (*Supplemental Fig. 1A-D*). However, TAs exposed to prolonged hypoxia exhibited a pronounced shift in fiber diameter towards smaller myofiber size, throughout the regenerative process (*Fig. 2B and 2C*). Morphometric quantification reveals a stronger difference of myofiber diameter at 14 and 28 dpi (*Fig. 2C*), at time points that mark the completion of muscle regeneration. This result correlates with significant reductions in regenerating TA cross-section area (*Fig. 2D*) and mass (*Fig. 2E*) in mice maintained under systemic hypoxia, both at 14 to 28 dpi. Since contralateral non-injured TAs did not show any sign of weight change under systemic hypoxia (*Fig. 2E*), this reduction in muscle mass observed in hypoxic-exposed regenerating TAs cannot be attributed to an indirect effect of systemic hypoxia, such as a decrease in mice motility or appetite. Surprisingly, in hypoxia-exposed regenerating TAs, the total number of myofibers per muscle was conserved (*Fig. 2F*) demonstrating that the hypotrophic fiber phenotype observed under systemic hypoxia is not associated with muscular hypoplasia.

**Figure 2:**
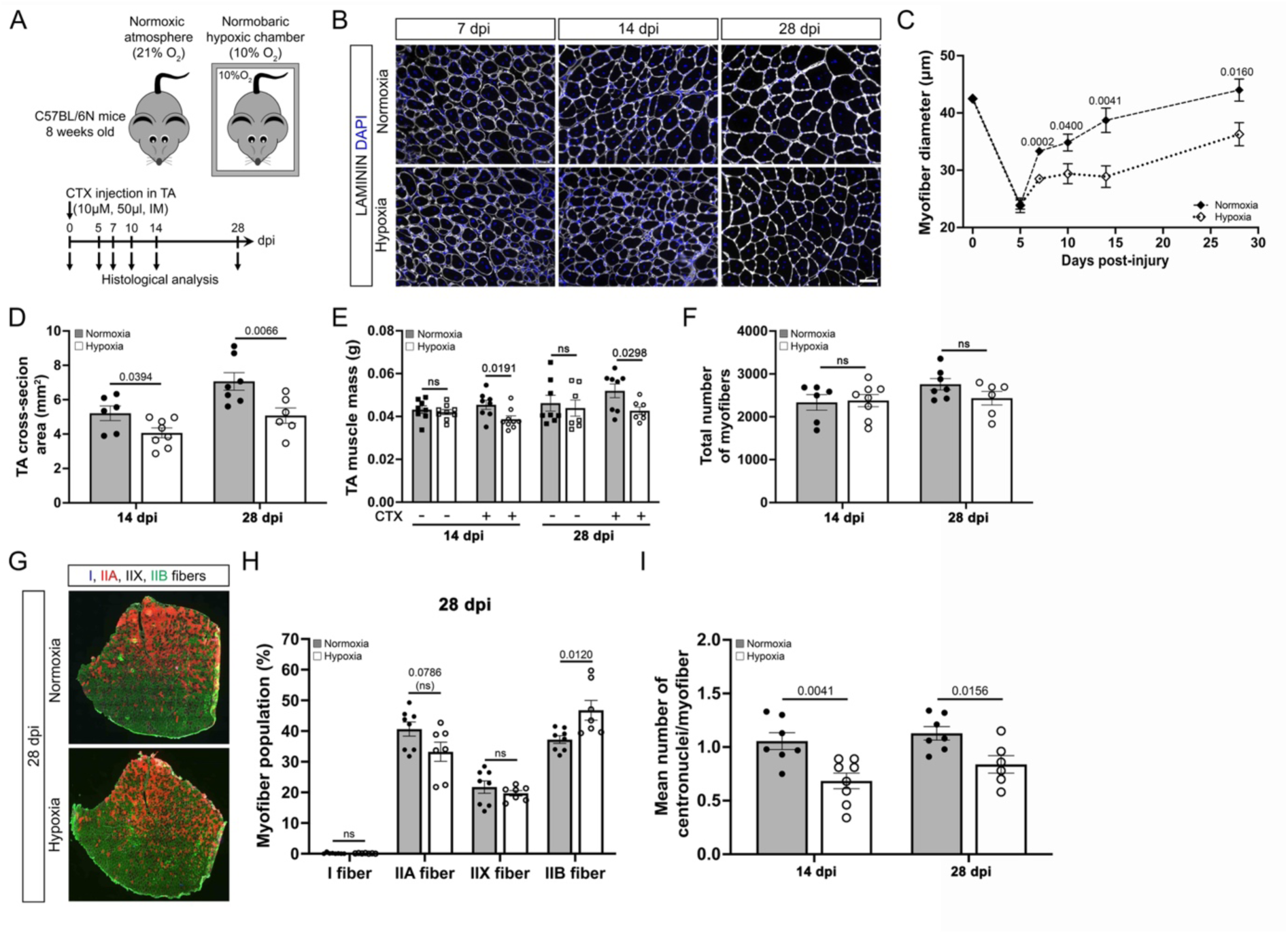
Blocking reoxygenation hampers muscle repair and leads to smaller regenerated myofibers, independent of any metabolic switch. **A.** Experimental design. Acute muscle injury was induced by cardiotoxin (CTX) intramuscular (IM) injection in the tibialis anterior (TA) of 8-week-old wild-type mice. Mice were then housed in standard normoxic atmosphere (21%O_2_) or in a normobaric hypoxic chamber (10%O_2_) from 0 to 28 days post-injury (dpi). **B.** Representative images of LAMININ^+^ myofiber (white) and nuclei (DAPI, blue) staining on cross-sections of CTX-injured TAs from mice exposed to normoxia or prolonged hypoxi, at 7, 14 and 28 dpi. Scale bar: 50μm. **C.** Quantification of myofiber diameters (μm) on cross-sections of CTX-injured TAs from mice exposed to normoxia or prolonged hypoxia, from 0 to 28 dpi. **D.** Quantification of CTX-injured TA cross-section areas of wild-type mice exposed to normoxia or prolonged hypoxia, at 14 and 28 dpi. **E.** Histograms showing muscle weights (g) of CTX-injured vs contralateral non-injured TAs from mice exposed normoxia or prolonged hypoxia, at 14 and 28 dpi. **F.** Quantification of the total number of myofibers in CTX-injured TAs from wild-type mice exposed to normoxia or prolonged hypoxia, at 14 and 28 dpi. **G.** Representative images of type-I (blue), type-IIA (red), type-IIB (green) myofiber and laminin (white) stainings performed on CTX-injured TA transversal muscle sections of wild-type mice exposed to normoxia or prolonged hypoxia, at 28 dpi. Scale bar 200μm. **H.** Quantification of myofiber type distribution (%) on CTX-injured TAs of wild-type mice exposed to normoxia or prolonged hypoxia, at 28 dpi. **I.** Quantification of the number of centronuclei per myofiber in CTX-injured TAs of wild-type mice exposed to normoxia (n = 6-7) or prolonged hypoxia (n = 6-8), at 14 and 28 dpi. **<u>Statistics</u>**: Results are expressed as means ± SEM. For **C, D, E, F, H and I.** Unpaired t-test. n=5-8 per time point.

To address whether this decrease in myofiber size was not the consequence of metabolic dysregulations, we evaluated the impact of prolonged hypoxia on myofiber protein synthesis and/or fiber typing in regenerating muscles. We first compared the *in vivo* protein synthesis rates in whole regenerating TAs using the puromycin incorporation based SUnSET assay, at 14 dpi (*Supplemental Fig. 1E*), as previously described^28^. Interestingly, we observed that the protein synthesis rates were similar in regenerating TAs exposed to prolonged hypoxia compared to normoxia (*Supplemental Fig. 1F and 1G*), demonstrating that blocking oxygen reoxygenation during muscle repair does not affect the hypertrophic growth of regenerating fibers. We next examined whether prolonged hypoxia affects the fiber type composition in the regenerating muscle. Fiber type analysis showed a lower percentage of type-IIA fibers and a higher percentage of type-IIB fibers in hypoxic-exposed regenerating muscles at 28 dpi (*Fig. 2G and 2H*). These data demonstrate that sustained hypoxia induces an oxidative-to-glycolytic transition of new-formed myofibers after an injury. Since glycolytic fibers are larger than oxidative fibers, it is unlikely that the glycolytic switch observed under hypoxic conditions is responsible for reducing muscle mass and myofiber size. Of interest, while multi-nucleated fibers were present abundantly in normoxia-exposed mice, the number of centralized nuclei per fiber was significantly decreased, both at 14 and 28 dpi, in hypoxia-exposed regenerating TAs (*Fig. 2I*), that could be the consequence of an altered proliferation, differentiation and/or fusion of myogenic cells.

Collectively, our results demonstrate the essential role of reoxygenation in adult muscle regeneration. Sustained hypoxia alters myogenesis and muscle repair, possibly through an impairment of myogenic cell differentiation and/or fusion rather than inducing a metabolic adaptation of myofibers to hypoxia.

### The dynamics of oxygen levels determine the fate of MuSCs during muscle repair

To test whether systemic hypoxia impacts MuSC number and specification *in vivo,* we first evaluated the basal level of oxygen specifically in this cell population following CTX-injury of TA muscle. Since MuSCs reached a peak of proliferation at 5 dpi (*Fig. 1D*), we performed co-immunostaining experiments against PAX7 and pimonidazole protein adducts at 0 and 5 dpi (*Fig. 3A*). At 0 dpi, the absence of pimonidazole staining suggests that quiescent PAX7^+^ cells are not in a hypoxic state in resting muscle (*Fig. 3B*). In contrast, all PAX7+cells in injured muscles acquire a hypoxic state at 5 dpi, concomitantly with their proliferation peak (*Fig. 3B*). We thus tested the influence of sustained hypoxia on MuSC fate *in vivo* following CTX-mediated injury in mice housed in a normal (21%O_2_) or hypoxic (10%O_2_ inhaled) normobaric atmosphere for 28 days (*Fig. 3C*). The increase in the number of PAX7^+^ cells at 5 dpi in CTX-injured TAs was not significantly different between mice exposed to hypoxia and control mice (*Fig. 3D and 3E).* However, under systemic hypoxia, PAX7^+^ cells accumulated in regenerating TAs after 7 and 10 dpi (*Fig. 3D and 3E*). To unravel this phenomenon, we decided to characterize the myogenic status of hypoxic-exposed PAX7+ cells at 7 dpi using PAX7, MYOD and KI67 triple immunostaining on injured TA muscle sections. As compared to normoxia, prolonged hypoxia showed an increased number of quiescent (PAX7^+^MYOD^-^KI67^-^) and proliferating (PAX7^+^KI67^+^) MuSCs along with a reduced number of differentiated (PAX7^-^MYOD^+^) myoblasts in regenerating muscles at 7 dpi (*Fig. 3F and 3G*). In contrast, systemic hypoxia did not affect the activation status of MuSCs (PAX7^+^MYOD^+^ cells) at 7 dpi (*Fig. 3F and 3G*). Together, these data demonstrate that suppressing reoxygenation *in vivo* by maintaining the mice under systemic hypoxia increases the self-renewal of MuSCs but limits their engagement towards differentiation, resulting in a transient accumulation of PAX7^+^ cells in hypoxic-exposed regenerating muscle.

**Figure 3:**
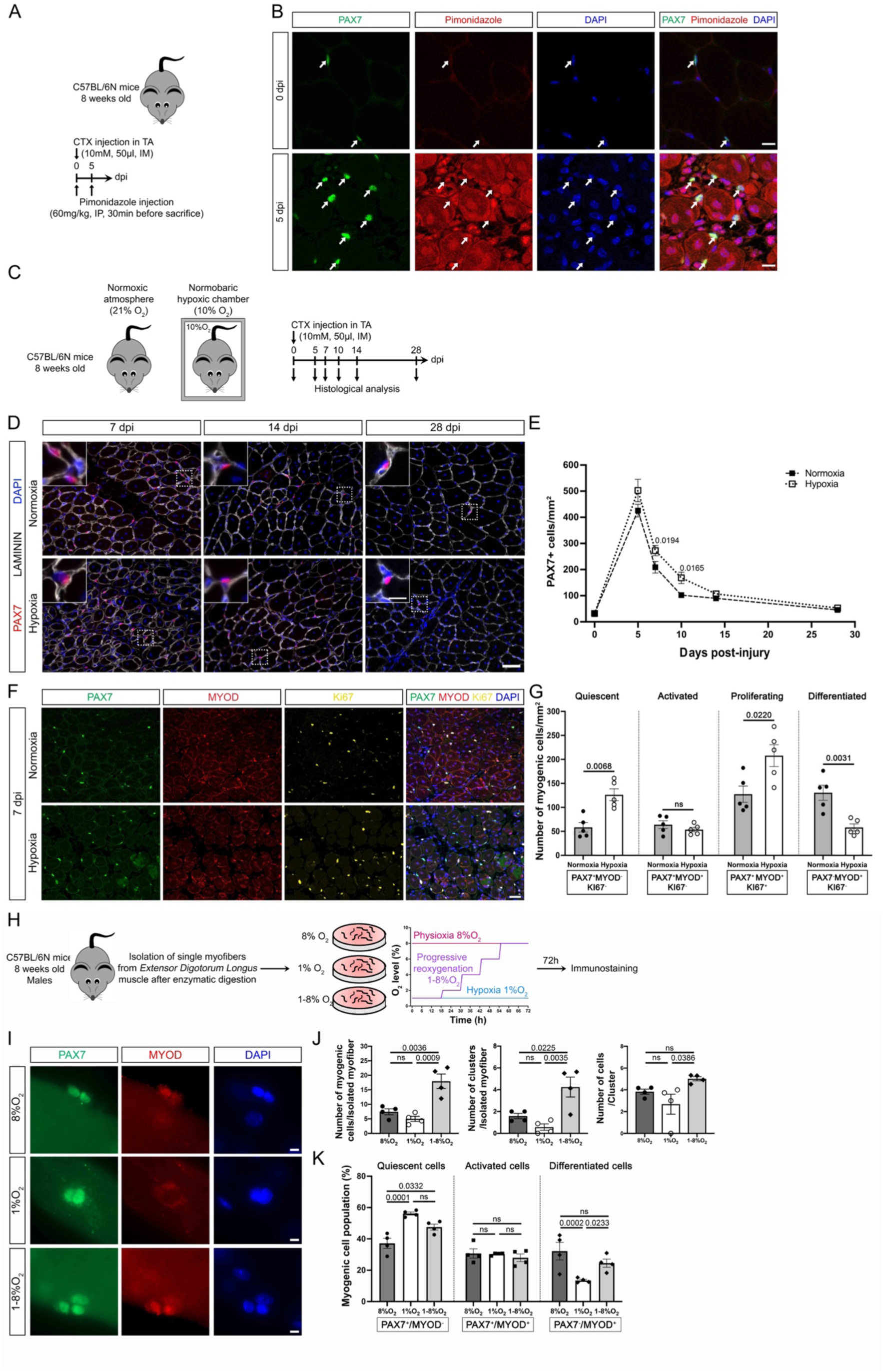
The dynamics of hypoxia phases in skeletal muscle defines the myogenic fate of muscle stem cells. **A.** Experimental design. Acute muscle injury was induced by cardiotoxin (CTX) intramuscular (IM) injection in the tibialis anterior (TA) of 8-week-old wild-type mice. Intraperitoneal (IP) injection of pimonidazole hypoxic probe were performed at 0 and 5 days post-injury (dpi). **B.** Representative co-immunofluorescence staining of pimonidazole hypoxic probe (red), PAX7^+^ cells (green) and nuclei (DAPI, blue) on CTX-injured TA cross-sections. White arrows show PAX7^+^ cells. Scale bar: 10 μm. **C.** Experimental design. Acute muscle injury was induced by CTX injection in the TA of 8-week-old wild-type mice. Mice were then housed in standard normoxic atmosphere (21%O_2_) or in a normobaric hypoxic chamber (10%O_2_), from 0 to 28 days post-injury (dpi). **D.** Representative pictures of PAX7^+^cells (red), LAMININ^+^ myofibers (white) and nuclei (DAPI, blue) on CTX-injured TA cross-sections of wild-type mice exposed to normoxia or prolonged hypoxia, at 7, 14 and 28 dpi. Scale bars: 50 μm (overviews) and 12.5 μm (insets). **E.** Quantification of PAX7^+^ cells/mm^2^ on CTX-injured TA cross-sections of wild-type mice exposed to normoxia or prolonged hypoxia. **F.** Representative co-immunofluorescence staining of PAX7 (red), MYOD (red), KI67 (yellow) and nuclei (DAPI, blue) on CTX-injured TA cross-sections of wild-type mice exposed to normoxia or prolonged hypoxia, at 7 dpi. Scale bar: 50 μm. **G.** Percentage of quiescent (PAX7^+^/MYOD^-^/KI67^-^), activated (PAX7^+^/MYOD^+^/KI67^-^), proliferating (PAX7^+^/MYOD^+^/KI67^+^) and differentiated (PAX7^-^/MYOD^+^/KI67^-^) cell populations on CTX-injured TA cross-sections of wild-type mice exposed to normoxia or prolonged hypoxia, at 7 dpi. **H.** Experimental design. Single myofibers were isolated from Extensor Digitorum Longus (EDL) muscles of 8-week-old wild-type mice and cultured *ex vivo* under physioxia (8%O_2_), prolonged hypoxia (1%O_2_) or transient hypoxia flowed by progressive reoxygenation (1-8%O_2_) during 72h. **I.** Representative pictures of PAX7 (green), MYOD (red) and nuclei (DAPI, blue) staining on isolated myofibers after 72h of culture under 8%, 1% or 1-8%O_2_. Scale bar: 20 μm. **J.** Quantification of the number of cells and clusters per myofiber and the number of cells per cluster on isolated myofibers after 72h of culture under 8%, 1% or 1-8%O_2_. **K.** Quantification of the percentage of quiescent PAX7^+^/MYOD^-^, activated PAX7^+^/MYOD^+^ and differentiated PAX7^-^/MYOD^+^ cells on isolated myofibers cultivated under 8%, 1% or 1-8%O_2_ for 72h. At 8%O_2_, 187 cells were counted on 25 myofibers; at 1%O_2_, 164 cells were counted on 32 myofibers; at 1-8%O_2_, 581 cells were counted on 32 myofibers. **<u>Statistics</u>:** Results are expressed as means ± SEM. For **E.** Unpaired t-test. n=5-8 per time point per group. For **G.** Unpaired t-test. n=5 per atmosphere exposure. For **J.** and **K.** One-way ANOVA followed by Tukey’s post-test. n=4 per O_2_ level.

To directly address the impact of hypoxia and/or reoxygenation on the status of MuSCs, we performed complementary experiments on *ex vivo* floating myofibers. Myofiber culture conditions allow MuSCs to become activated, start proliferating into cluster (24-48h), and proceed to myogenic differentiation or self-renewal of the quiescent pool at 72h of culture^7^. Single myofiber isolated from EDL muscles were maintained for 72h, either in physioxia (8%O_2_), in sustained hypoxia (1%O_2_) and in transient hypoxia followed by progressive reoxygenation (1% to 8%O_2_) (*Fig. 3H*). As compared to physioxia, sustained hypoxic culture conditions had no significant impact on the overall growth of MuSCs at 72h (*Fig. 3I and 3J).* However, prolonged hypoxia decreased the ratio of differentiating Pax7^-^/MYOD^+^ cells towards an increase of Pax7^+^MYOD^-^ self-renewing cells (*Fig. 3K)*. Strikingly, transient hypoxia followed by progressive reoxygenation potentiated myogenic cell expansion and cluster density, as compared to prolonged hypoxic or physioxic culture conditions (*Fig. 3I and 3J)*. Moreover, although the capacity of MuSCs to self-renew *in vitro* was less under transient hypoxia/reoxygenation than prolonged hypoxia, the differentiation capacity of PAX7+ cells was restored under the reoxygenation process (*Fig. 3K)*. Both prolonged hypoxia and transient hypoxia/reoxygenation did not modulate the activation status of MuSCs, as observed *in vivo* (*Fig. 3G*). Altogether, our data suggest that blocking muscle reoxygenation through prolonged hypoxia directly disrupts the myogenic fate of MuSCs, by enhancing their self-renewal at the expense of their differentiation.

### Hypoxia alters myoblastic fusion capacity of MuSCs independently of their commitment towards differentiation

We next addressed whether prolonged hypoxia affects myoblast cell fusion to regenerating myofibers by impairing MuSC differentiation potential or by directly impacting the fusion capacity of myogenic cells. For this, we cultured FACS-sorted GFP^+^ primary MuSCs, which spontaneously fuse *in vitro* in normoxic atmosphere after 72h (*Fig. 4A*). Using MHC immunostaining, we showed that MuSCs maintained under prolonged hypoxia form less and smaller myotubes compared to physioxic culture conditions (*Fig. 4B and 4C)*. In contrast, transient hypoxia followed by progressive reoxygenation rescued the fusion capacity of MuSCs (*Fig. 4B and 4C*).

**Figure 4:**
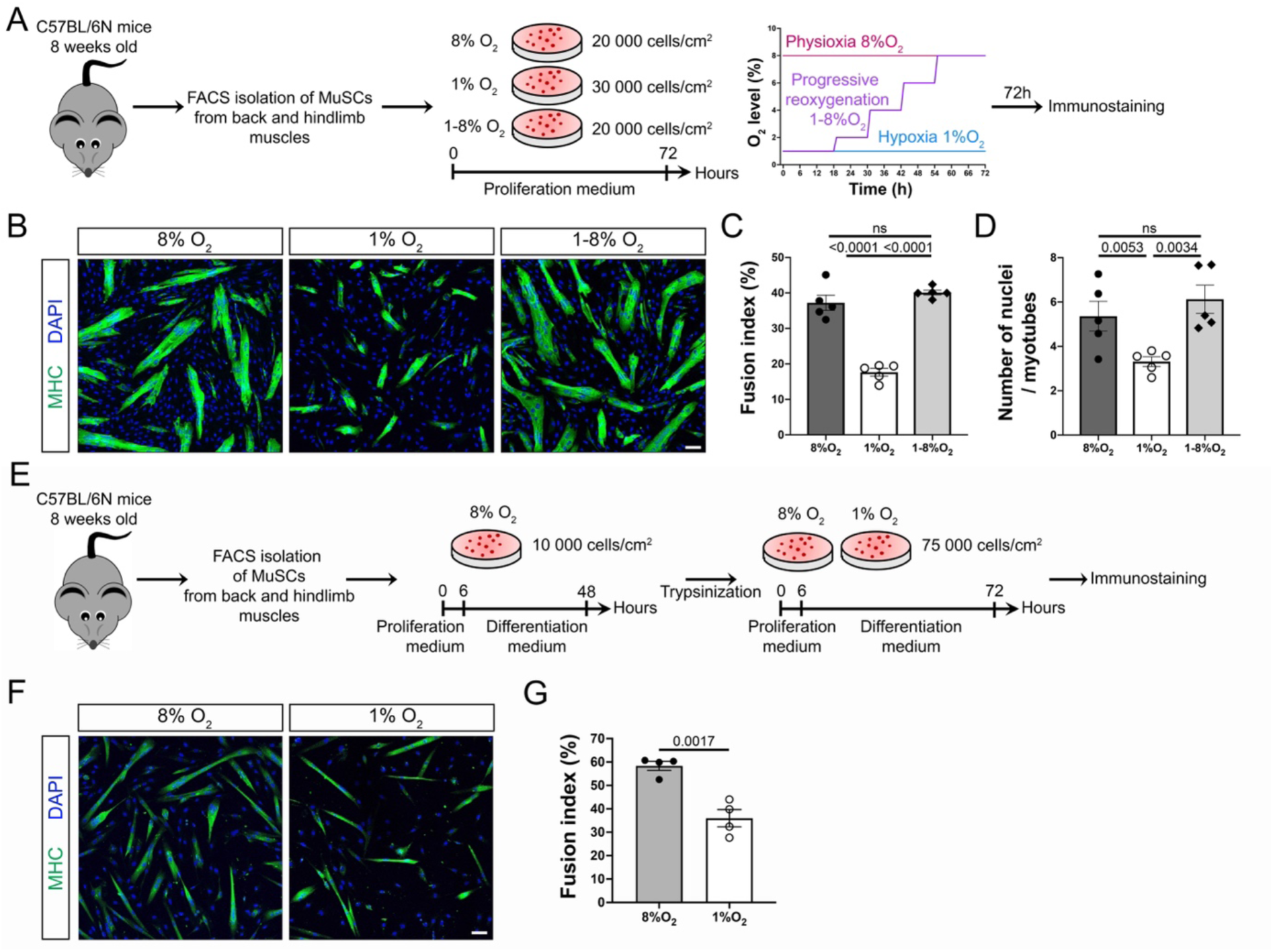
Hypoxia impairs the fusion capacity of murine muscle stem cells. **A.** Experimental design. FACS-sorted MuSCs were isolated from wild-type mice and plated with appropriate densities to ensure same confluence after 72h of culture under physioxia (8%O_2_), prolonged hypoxia (1%O_2_) and transient hypoxia followed by progressive reoxygenation (1-8%O_2_). **B.** Representative pictures of MHC^+^ (Myosin Heavy Chain) myotubes and nuclei (DAPI, blue) staining after 72h of myogenic cell culture under physioxia (8%O_2_), hypoxia (1%O_2_) and progressive reoxygenation (1-8%O_2_). Scale bar: 50 μm. **C.** Quantification of fusion index (number of nuclei per myotube divided by total number of nuclei, %) and of the number of nuclei per myotubes after 72h of myogenic cell culture under 8%, 1%O_2_ or 1-8%O_2_. **D.** Experimental design. FACS-sorted MuSCs were isolated from wild-type mice and plated at low density for 48h under 8%O_2_ to allow the synchronization of myoblasts, as myocytes. After trypsinization, myocytes were seeded at high density to allow their fusion after 72h of culture under physioxia (8%O_2_) or prolonged hypoxia (1%O_2_). **E.** Representative pictures of MCH^+^ myotubes and nuclei (DAPI, blue) staining after 48h of culture under 8%O_2_ followed by 72h of culture under 8% or 1%O_2_. Scale bar: 50 μm. **F.** Quantification of fusion index (%) of synchronized myoblasts after 72h of culture under 8% or 1%O_2_. **<u>Statistics</u>:** Results are expressed as means ± SEM. For **C.** One-way ANOVA followed by Tukey’s post-test. n=5 per O_2_ level. For **F.** Unpaired t-test. n=4 per O_2_ level.

As prolonged hypoxia reduced the differentiation status of MuSCs (*Fig. 3G and 3K*), it may be a confounding factor *in vitro* to access the direct impact of hypoxic conditions on myoblast fusion potential. To address this question, we seeded PAX7+ cells at low density and directly cultivated them in differentiation medium, allowing the synchronization of myocytes (*Fig. 4E*). Cells engaged in differentiation were then seeded at high density to allow their fusion (*Fig. 4E*). We then quantified the fusion potential of primary myoblasts in high-density culture, in both physioxic and prolonged hypoxic conditions *(Fig.4F*). Using this strategy, we conclude that prolonged hypoxia directly affects the fusion potential of MuSCs, as compared to physioxic conditions, independently of their differentiation status (*Fig. 4E and 4F*).

### Prolonged hypoxia impairs skeletal muscle repair by limiting the differentiation and fusion capacity of MuSCs in HIF-1α-independent manner

Overall, our results highlight the importance of transient hypoxia followed by progressive reoxygenation process in preserving the efficient myogenic differentiation and fusion abilities of MuSCs for proper skeletal muscle repair. However, the molecular mechanism by which prolonged hypoxia impairs skeletal muscle repair remains to be elucidated. In many cells, the cellular adaptation to hypoxic stress is orchestrated through the activation of the HIF-1 signaling pathway^10^. To evaluate whether blocking progressive reoxygenation by chronic hypoxia modulates the differentiation and fusion potential of MuSCs through HIF-1α signaling, we generated a MuSC-specific HIF-1α conditional knockout mouse model (HIF cKO) by combining *Pax7^CreERT^*^2^*^/+^* allele with *HIF-1α^fl/lf^* (*Supplemental Fig. 2A*). We validated the efficiency of HIF-1α deletion following tamoxifen-induced Cre ablation of HIF-1α in HIF cKO MuSCs compared to HIF CTRL MuSCs at the mRNA (*Supplemental Fig. 2A and 2B*) and protein levels (*Supplemental Fig. 2C, 2D and 2E*). We then evaluated the specific role of HIF-1α on MuSC function in HIF cKO and CTRL mice exposed to normoxia (21% O_2_) or systemic hypoxia (10%O_2_ inhaled) during skeletal muscle regeneration (*Fig. 5A*). In normoxic atmosphere, HIF cKO mice displayed similar regenerative dynamics of myofiber growth (*Fig. 5B and 5C*), muscle size (*Fig. 5C*), myogenic cell expansion (*Supplemental Fig. 3A-C*) and fiber type proportions (*Supplemental Fig. 3D and 3E*) compared with HIF CTRL mice, at 7 or 28 dpi. Similarly, HIF cKO mice maintained in prolonged hypoxia show similar accumulation of PAX7^+^ cells at 7 dpi (*Supplemental Fig. 3B and 3C*), oxidative to glycolytic fiber-type switch (*Supplemental Fig. 3D and 3E)* and muscle atrophy (*Fig. 5B and 5C*) than HIF CTRL mice at 28 dpi. These data demonstrate that HIF-1α expression in MuSCs is dispensable for muscle regeneration, regardless of the dynamics of oxygen level. To further explore the role of HIF-1α in the myogenic potential of MuSCs, we performed complementary experiments on floating myofibers isolated from HIF cKO and HIF CTRL EDL muscles (*Fig. 5D*). Using these experiments, we confirm that HIF-1α depletion had no impact on the differentiation state of MuSCs, whatever the oxygen level used (*Fig. 5E and 5F*). Moreover, using FACS-isolated MuSCs from HIF cKO and HIF CTRL muscles, we showed that the reduction of myoblast fusion capacity upon prolonged hypoxia was independent of HIF-1α expression (*Fig. 5.G-I*). Of interest, we finally demonstrate that the enhancement of MuSC self-renewal under sustained hypoxia was abrogated towards increased MuSC activation when HIF-1α was depleted in MuSCs (*Fig. 5E and 5F*).

**Figure 5:**
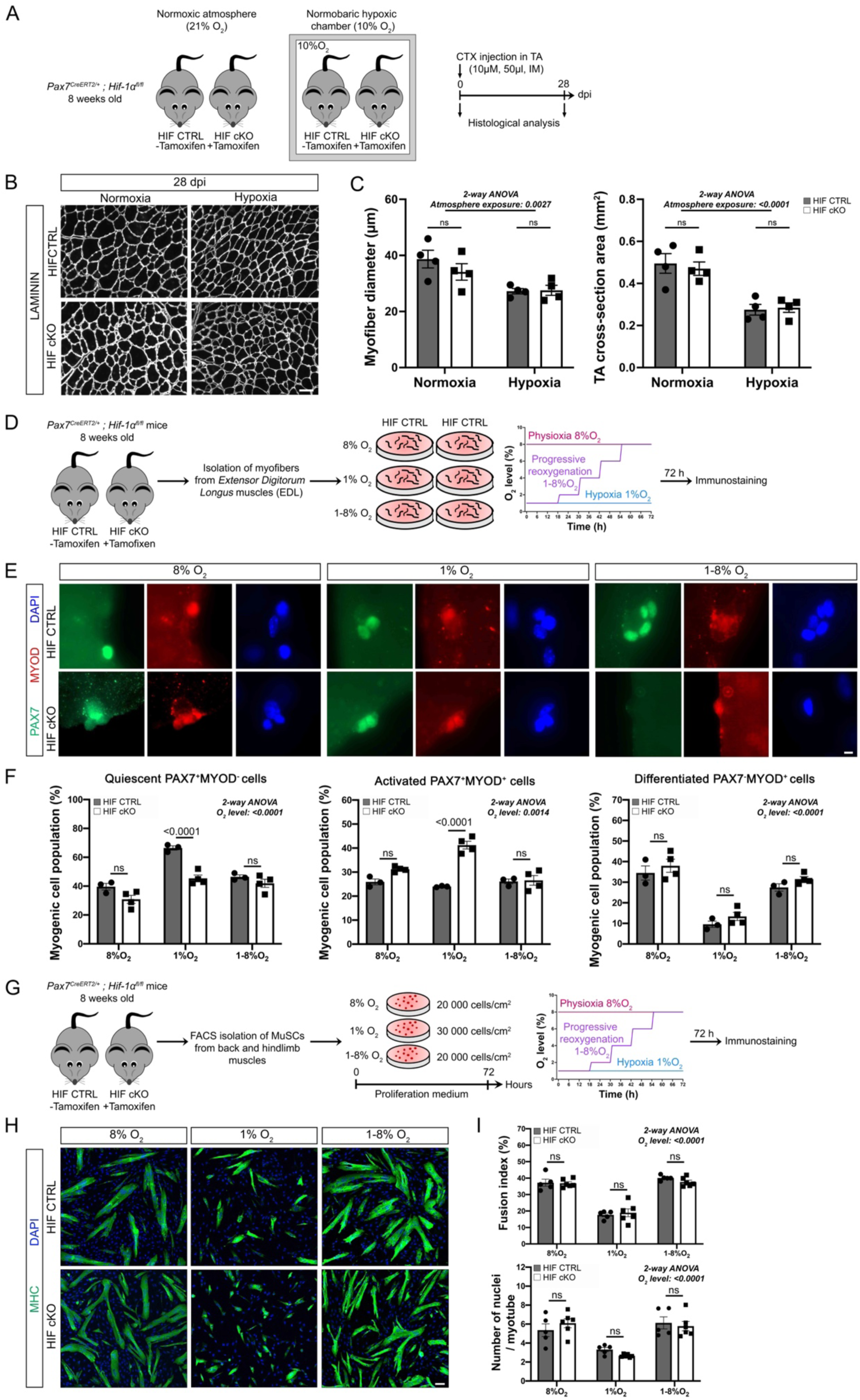
Impact of conditional deletion of HIF-1α in muscle stem cells on muscle repair and myogenic program. **A.** Experimental design. Acute muscle injury was induced by cardiotoxin (CTX) intramuscular (IM) injection in tibialis anterior (TA) muscles from 8-week-old HIF CTRL (Pax7^CreERT2/+^; HIF-1α^fl/fl^ without tamoxifen) and HIF cKO (Pax7^CreERT2/+^; HIF-1α^fl/fl^ with Cre recombinase induction by tamoxifen) mice. Mice were then housed in standard normoxic atmosphere (21%O_2_) or in a normobaric hypoxic chamber (10%O_2_), from 0 to 28 days post-injury (dpi). **B.** Representative pictures of LAMININ (white) staining on CTX-injured TA cross-sections from HIF CTRL and HIF cKO mice exposed to normoxia or prolonged hypoxia, at 28 dpi. Scale bar: 50μm. **C.** Quantification of the myofiber diameter (μm) and the number of myofiber per mm^2^ of CTX-injured TA cross-sections from HIF CTRL and HIF cKO mice exposed to normoxia or prolonged hypoxia, at 28 dpi. **D.** Experimental design. Single myofibers were isolated from Extensor Digitorum Longus (EDL) muscles of 8-week-old HIF CTRL or HIF cKO mice and cultured *ex vivo* under physioxia (8%O_2_), prolonged hypoxia (1%O_2_) or transient hypoxia followed by progressive reoxygenation (1-8%O_2_) during 72h. **E.** Representative staining of PAX7 (green), MYOD (red) and nuclei (DAPI, blue) on isolated myofibers from HIF CTRL or HIF cKO after 72h of culture under 8%, 1%O_2_ or 1-8%O_2_. Scale bar: 20 μm. **F.** Percentage of quiescent (PAX7^+^/MYOD^-^), activated (PAX7^+^/MYOD^+^) and differentiated (PAX7^-^/MYOD^+^) cell populations obtained on isolated myofibers from EDL of HIF CTRL or HIF cKO mice, after 72h of culture under 8%O_2_, 1%O_2_ or 1-8%O_2_. For HIF CTRL, n=3 with a counting of 654 cells on 38 myofibers at 8%O_2_, 337 cells on 43 myofibers at 1%O_2_ and 1076 cells on 49 myofibers at 1-8%O_2_. For HIF cKO, n=4 with a counting of 643 cells on 47 myofibers at 8%O_2_, 292 cells on 57 myofibers at 1%O_2_ and 788 cells on 62 myofibers at 1-8%O_2_. **G.** Experience design. FACS-sorted MuSCs were isolated from HIF CTRL or HIF cKO mice and plated with appropriate densities to ensure same confluence after 72h of culture under physioxia (8%O_2_), prolonged hypoxia (1%O_2_) and transient hypoxia followed by progressive reoxygenation (1-8%O_2_). **H.** Representative pictures of MHC^+^ (Myosin Heavy Chain) myotubes and nuclei (DAPI, blue) staining after 72h of culture under 8%, 1% or 1-8% O_2_. Scale bar: 50 μm. **I.** Evaluation of fusion index (%) and number of nuclei per myotube after 72h of culture under 8%, 1% or 1-8% O_2_. **<u>Statistics</u>:** Results are expressed as means ± SEM. For **C, F and I.** 2-way ANOVA followed by Sidak’s post-tests. n=4 per mice group per atmosphere exposure (C), n=3-4 per mice group per O_2_ level (F) and n=5-6 per mice group per per O_2_ level (I).

Collectively, these results demonstrate that reducing reoxygenation by maintaining the mice under systemic hypoxia impairs skeletal muscle repair by limiting the differentiation and fusion capacity of MuSCs in HIF-1α-independent manner, while it favors their return into quiescence through HIF-1α activation.

### REV-ERBα regulates the differentiation and fusion potential of MuSCs under prolonged hypoxia

We demonstrated that the impairment of MuSC differentiation and fusion capacity under prolonged hypoxia is not regulated by HIF-1α. To decipher which molecular mechanisms could explain this phenotype, we performed a bulk-RNAseq on freshly isolated GFP+ myogenic cells from 7 dpi TAs isolated from mice exposed or not to a hypoxic atmosphere (*Fig. 6A*). Among the 22373 detected genes, 3728 genes were differentially expressed (DEGs) under prolonged hypoxia, with 2022 downregulated genes and 1706 upregulated genes compared to normoxic atmosphere (pvalue ≤ 0.05 ; *Fig. 6B*). Among those DEGs, the quiescence marker, *Pax7* was increased while *Myod* and *Myog,* two myogenic differentiation factors and the fusion marker, *Myomaker* (*Mymk*) were decreased in hypoxic-exposed FACS-sorted PAX7+ cells (*Fig. 6C*). These results confirm our *in vivo* and *ex vivo* data demonstrating that blocking reoxygenation after transient hypoxia facilitates MuSC return into quiescence but blunts their differentiation and fusion capacities.

**Figure 6:**
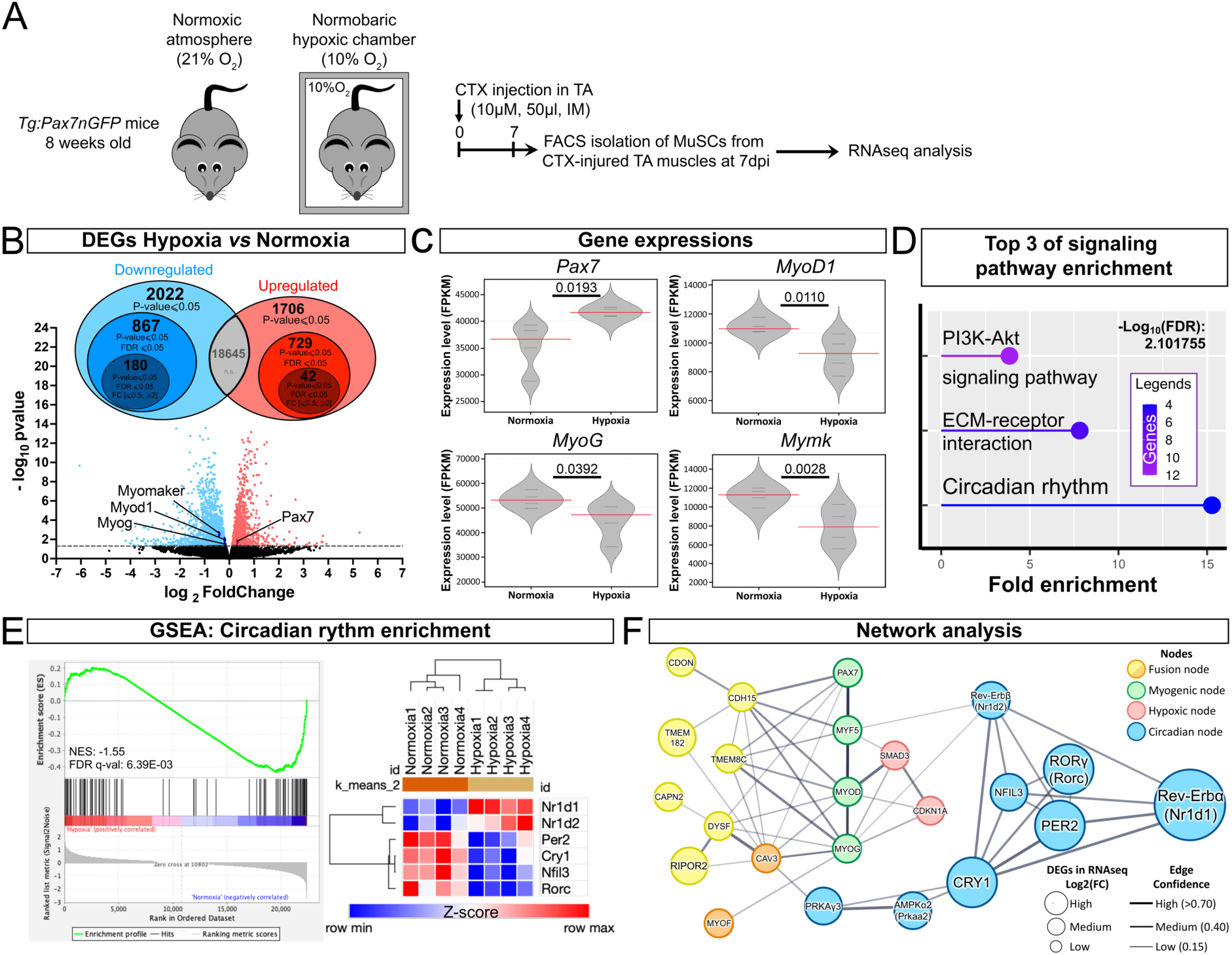
Transcriptomic analysis of PAX7^+^ cells isolated from injured *Tibialis Anterior* muscles of mice exposed to hypoxic *versus* normoxic atmosphere. **A.** Experimental design. Acute muscle injury was induced by cardiotoxin (CTX) intramuscular (IM) injection in the tibialis anterior (TA) of 8-week-old Tg:Pax7nGFP mice. Mice were then housed in standard normoxic atmosphere (21%O_2_) or in a normobaric hypoxic chamber (10%O_2_), from 0 to 7 days post-injury (dpi). FACS-sorted PAX7+ (GFP+) cells were isolated from CTX-injured TAs at 7 dpi to perform a bulk RNAseq analysis. n=4 mice per atmosphere exposure. **B.** Differential Gene Expression (DEG) of PAX7+ cells isolated from injured TAs of mice exposed to hypoxic versus normoxic atmosphere. Volcano plot was obtained from p-values > 0.05. **C.** Violin plots representing myogenic quiescence (*Pax7*), differentiation (*MyoD1, MyoG*) and fusion (*Mymk* = myomaker) genes significatively modulated in PAX7+ cells exposed to hypoxia. **D.** Top 3 of signalling pathways obtained from KEGG enrichment analysis of all DEGs with FDR (False Discovery Rate) ≤ 0.05 and fold-change [0.5 ≤ ; ≥ 2]. **E.** Gene set enrichment analysis (GSEA) of circadian rhythm pathway and Heatmap showing the main 6 genes actors of circadian rhythm significatively (FDR ≤ 0.05 and fold-change [0.5 ≤ ; ≥ 2]) upregulated (red) or down-regulated (blue) in our RNAseq. **F.** Network representation of the protein-protein interactions of genes differentially regulated in PAX7+ cells exposed to hypoxia with FDR ≤ 0.05, obtained by using STRING.

We then selected the 222 highest modulated DEGs under prolonged hypoxia with FDR ≤ 0.05 and fold-change [0.5 ≤ ; ≥ 2] to perform a KEGG pathway enrichment. Among the top-3 enriched pathways was the PI3K-Akt signalling pathway (*Fig. 6D*), with numerous genes known to control muscle stem cell fate and repair under normoxic^29,30^ or hypoxic conditions^3^. Another enriched signalling pathway implicated extracellular matrix-receptor interaction genes (*Fig. 6D*), that are critical regulators of muscle stem cell function, skeletal muscle development and repair^30^. Surprisingly, the most top-enriched pathway in our data set was the circadian rhythm which revealed a small but highly modulated number of DEGs (*Fig. 6D*). To confirm and assess which circadian rhythm genes were modulated under prolonged hypoxia, we realized a Gene Set Enrichment Analysis (GSEA) for this specific pathway (*Fig. 6E*). In 7 dpi injured TAs, we confirmed that MuSCs isolated from mice maintained under systemic hypoxia presented a significant modulation of genes encoding the core regulators of the circadian rhythm. Indeed, under prolonged hypoxia, circadian clock activators, such as *Cry1*, *Per2*, *Nfil3* and *Rorc* were downregulated while circadian clock repressors, such as *Nr1d1* and *Nr1d2* (coding Rev-ERBα and Rev-ERBβ, respectively) were upregulated in PAX7+ cells (*Fig. 6E*).

To decipher the molecular interplay more deeply between the circadian rhythm and muscle repair under prolonged hypoxia, we generated a computational model of protein-protein interactions networks (PPIs) using STRING v.11 plugins (http://string-db.org). We visualized on Cytoscape software both physical and functional interaction between these DEGs for 3 clusters specifically targeting the myogenic fusion node, the core circadian clock node and the hypoxia-mediated myogenesis node. The list of genes involved in these 3 specific clusters were retrieved from GeneOntology, KEGG and Wikipathways (WP5023, 5024, 5025), respectively. According to this predictive network analysis, the edges between hypoxia and myogenesis nodes can directly explain the impact of prolonged hypoxia on the myogenic status of MuSC and its consequence on muscle repair (*Fig. 6F*). On the other hand, this projective network approach identifies the circadian rhythm as a key regulator of the myogenic program, either by the upstream modulation of myogenic differentiation factors or directly by affecting late myoblastic fusion (*Fig. 6F*). The highest regulated circadian clock DEGs is the circadian clock repressor, *Nr1d1* (Rev-ERB) which was markedly increased by more than 2.8-fold in hypoxia-exposed PAX7+ cells, when muscle reoxygenation was blocked. Using freshly isolated GFP+ myogenic cells from 7 dpi TAs taken from mice exposed or not to a hypoxic atmosphere (*Fig. 7A*), we confirmed the upregulation of *Nr1d1* and the downregulation of *Cry1* and *Per2* under systemic hypoxia (*Fig. 7B*).

**Figure 7:**
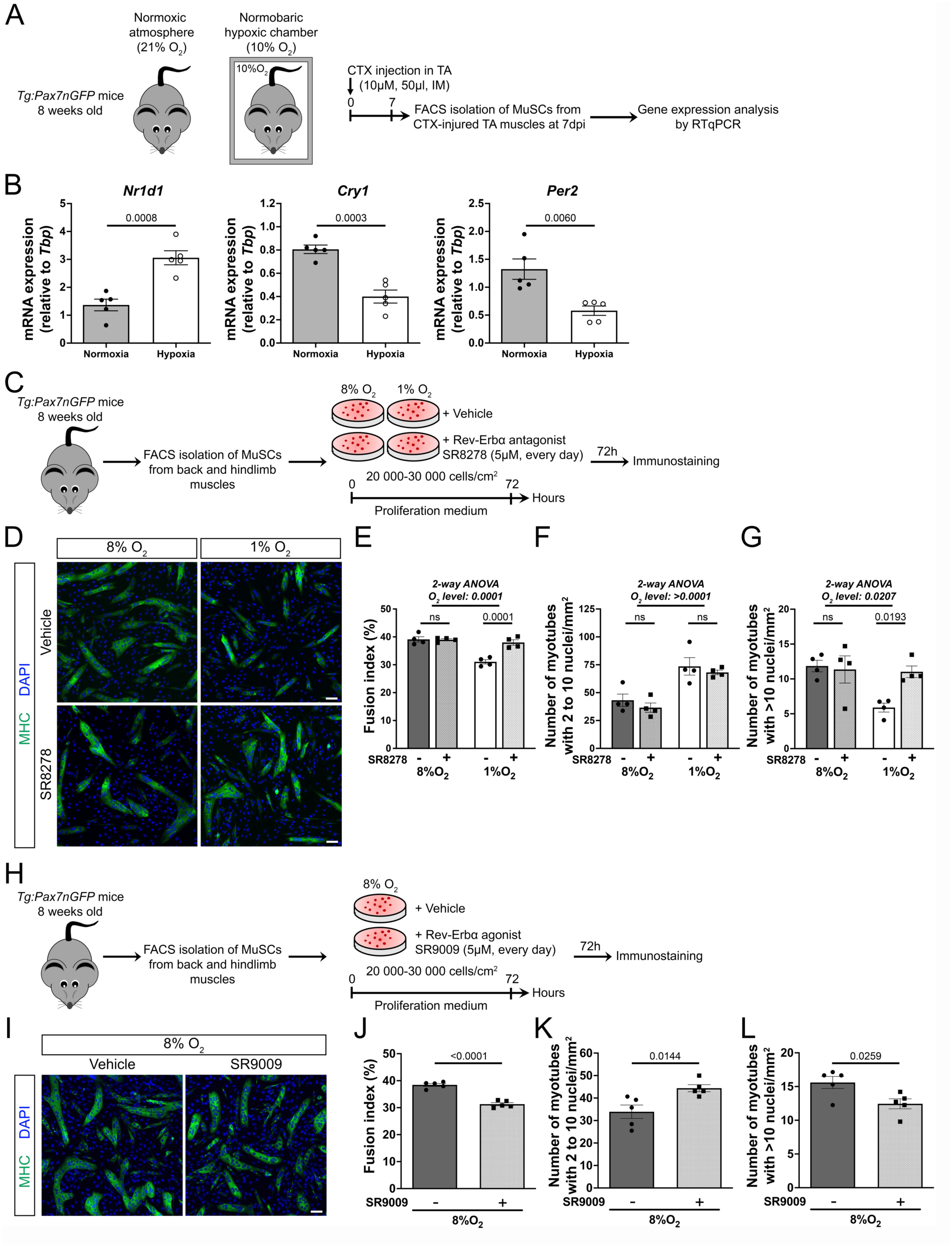
Upregulation of *Nr1d1* (Rev-ERBα) by hypoxia controls myogenic fusion. **A.** Experimental design. Acute muscle injury was induced by cardiotoxin (CTX) intramuscular (IM) injection in the tibialis anterior (TA) of 8-week-old Tg:Pax7nGFP mice. Mice were then housed in standard normoxic atmosphere (21%O_2_) or in a normobaric hypoxic chamber (10%O_2_), from 0 to 7 days post-injury (dpi). FACS-sorted PAX7+ (GFP+) cells were isolated from CTX-injured TAs at 7 dpi to perform a gene expression analysis. **B.** Histograms showing the relative expression of circadian clock effectors *Nr1d1 (*nuclear receptor subfamily 1 group D member 1), *Cry1* (cryptochrome circadian regulator 1) and *Per2* (period circadian regulator 2) normalized to *Tbp* (TATA-box Binding Protein) on FACS-sorted PAX7+ (GFP+) cells isolated from CTX-injured TAs at 7 dpi. **C.** Experimental design. FACS-sorted PAX7+ (GFP+) cells were isolated from 8-week-old Tg:Pax7nGFP mice and plated with appropriate densities to ensure same confluence after 72h of culture under physioxia (8%O_2_) or prolonged hypoxia (1%O_2_), in the presence of the Rev-ERBα antagonist, SR8278 or its vehicle. **D.** Representative pictures of MHC^+^ (Myosin Heavy Chain) myotubes and nuclei (DAPI, blue) staining after 72h of culture under 8% or 1%O_2_, with the Rev-ERBα antagonist SR8278 or its vehicle. Scale bar: 50 μm. **E.** Evaluation of fusion index (%) after 72h of culture under 8% or 1%O_2_, with the Rev-erbα antagonist SR8278 or its vehicle. **F.** Quantification of the number of small myotubes containing between 2 and 10 nuclei/mm^2^, after 72h of myogenic myogenic cell culture under physioxia (8%O_2_) or hypoxia (1%O_2_), with the Rev-erbα antagonist SR8278 or its vehicle. **G.** Quantification of the number of large myotubes containing more than 10 nuclei/mm^2^, after 72h of culture under 8% or 1%O_2_, with the Rev-erbα antagonist SR8278 or its vehicle. **H.** Experimental design. FACS-sorted PAX7+ (GFP+) cells were isolated from 8-week-old Tg:Pax7nGFP mice and plated with appropriate densities to ensure same confluence after 72h of culture under physioxia (8%O_2_) in the presence of the Rev-ERBα agonist, SR9009 or its vehicle. **I.** Representative pictures of MHC^+^ (Myosin Heavy Chain) myotubes and nuclei (DAPI, blue) staining after 72h of culture under 8%O_2_, with the Rev-ERBα agonist SR9009 or its vehicle. Scale bar: 50 μm. **J.** Evaluation of fusion index (%) after 72h of culture under 8% with the Rev-erbα agonist SR9009 or its vehicle. **F.** Quantification of the number of small myotubes containing between 2 and 10 nuclei/mm^2^, after 72h of myogenic myogenic cell culture under physioxia (8%O_2_) with the Rev-erbα agonist SR9009 or its vehicle. **G.** Quantification of the number of large myotubes containing more than 10 nuclei/mm^2^, after 72h of culture under 8% with the Rev-erbα agonist SR9009 or its vehicle. **<u>Statistics</u>:** Results are expressed as means ± SEM. For **E, F and G.** 2-way ANOVA followed by Sidàk’s post-tests. n=4 per O_2_ level and per condition. For **J, K and L.** Unpaired t-test. n=5 per condition.

Our results suggest that hypoxia-circadian clock interactions may reduce myogenesis though the regulation of Rev-ERBα in MuSCs. To test our predictive model, we performed *in vitro* experiments on FACS-isolated MuSCs cultured under physioxia (8%O_2_) or hypoxia (1%O_2_) in the presence or not of a Rev-ERBα antagonist, SR8278 (*Fig. 7C*). In physioxia, inhibiting Rev-ERBα *in vitro* had no influence on the number or the fusion capacity of MuSCs, indicating that Rev-ERBα does not affect MuSC fate under normoxia (*Fig. 7D-G*). In contrast, the decreased fusion index of myogenic cells under prolonged hypoxia was fully restored when Rev-ERBα was blocked (*Fig. 7D-G*). This preservation of myogenic fusion potential in the presence of Rev-ERBα inhibitor was associated with the specific increase in the number of large mature myotubes (>10 nuclei; *Fig. 6K*), indicating that Rev-ERBα mostly impacts the late steps of myogenic fusion process under prolonged hypoxia, notably by decreasing myonuclear accretion. We then performed the reverse experiment by incubating freshly-isolated MuSCs in the presence of a specific Rev-ERBα agonist, SR9009 under physioxia (8%O_2_), to mimic the increase of Rev-ERBa under prolonged hypoxia (*Fig. 7H*). Of interest, incubation of physioxic-exposed myogenic cells with SR9009 impairs the fusion capacity of MuSCs (Fig. *7I-L*) and decreases the number of large mature myotubes (>10 nuclei; *Fig. 7L*), reproducing the inhibitory impact of hypoxia on myogenic cell fusion. Altogether, our results demonstrate that blocking reoxygenation through sustained hypoxia impairs myogenic cell fusion through Rev-ERBα upregulation.

## DISCUSSION

Altogether our results demonstrate that progressive reoxygenation after transient hypoxia is essential to preserve the capacities of MuSCs to differentiate and fuse and to properly repair the damaged muscle. We further show that blocking reoxygenation by prolonged hypoxia impairs the myogenic program in a HIF-1α-independent manner but though the upregulation of Rev-ERBα.

Our work confirms that efficient skeletal muscle repair correlates with myo-angiogenesis coupling, as already shown in the litterature^9^. However, our study is the first to decipher the dynamics of muscle oxygenation throughout the muscle repair process. We show that after acute injury, muscle repair is initiated in a low oxygen environment followed by progressive reoxygenation. In the regenerating muscle, variation of oxygen levels synchronizes with muscle revascularization. This result agrees with recent works showing that muscles are not hypoxic in the steady state but become hypoxic soon after the injury^32^. We further show that transient hypoxia following muscle injury correlates with the induction of the HIF signaling, as reported previously^12^. Hence, we show that oxygen level in muscle tissue inversely correlates with the mobilization of MuSCs during muscle repair. Indeed, while in the resting muscle, quiescent MuSCs reside in a normoxic environment, proliferating PAX7^+^ cells are in a hypoxic state after 5 dpi in the TA-injured muscle. This result is in sharp contrast with those of Xie and colleagues who claimed that quiescent MuSCs in resting muscle are intriguingly inherently hypoxic, as they expressed high level of HIF-2α at the surface of isolated myofibers^33^. This affirmation is rather surprising given that quiescent MuSCs are located in the immediate vicinity of blood capillaries^34^, questioning their actual hypoxic state in healthy muscle. Although it was not address in their study, cellular stabilization of HIF-2α in quiescent MuSCs may implicate hypoxia-independent mechanisms, as it has been shown in numerous other settings^35^. Besides, no study has yet addressed the direct impact of intramuscular oxygen levels in regulating MuSC fate and muscle repair during the regeneration process.

Of interest, we show that reoxygenation of skeletal muscle after transient tissue hypoxia is essential for proper muscle repair. Indeed, maintaining mice in a hypoxic atmosphere uncouples myo-angiogenesis affecting muscle repair, and leads to a hypotrophic muscle phenotype, mimicking muscle atrophy observed in Highlanders, climbers or Chronic Obstructive Pulmonary Disease (COPD) patients exposed to long-lasting systemic hypoxia^36^. But as opposed to long-lasting hypoxia, the hypotrophic phenotype observed under prolonged hypoxia in our study is restricted to the regenerating muscle whereas the contralateral uninjured muscle remains unaffected. This result confirms that in our case, systemic hypoxia specifically impedes the muscle repair process by blocking intramuscular reoxygenation.

Whether the decrease of myofiber size could be due to a critical protein loss^37^, we invalidate this hypothesis as we didn’t observe any change in protein synthesis rates. In the same way, this presence of smaller myofibers under prolonged hypoxia could not be explain by the glycolytic fiber-type switch from type-IIA to type-IIB, as observed in pathological context of COPD^38^ or peripheral arterial disease (PAD)^39,40^, since type-IIB are the largest myofibers in mice^41^. These findings, combined with the fact that hypotrophic regenerating myofibers present a reduced number of myonuclei argue for a deregulation of MuSC myogenic abilities upon prolonged hypoxia.

In the literature, the impact of hypoxia on the fate of MuSCs has been mainly evaluated by *in vitro* studies and has led to controversial results^42^, in particular concerning the effect of hypoxia on MuSC proliferation. Indeed, some studies show that hypoxia promotes myogenic proliferation^43,44^ while some others demonstrate the opposite^24,45,46^. Such a discrepancy could be attributed to the different methods used to induce hypoxia, the various degree of oxygen level tested (from 0,01% to 6%O_2_), multiple cell types from different species and the diverse methods to quantify proliferation. Our work show that sustained hypoxia (1%O_2_) as compared to physioxia (8%O_2_) tends to decrease the expansion of MuSCs *ex vivo* on floating myofibers. In contrast, transient hypoxia followed by progressive reoxygenation enhances myogenic cell expansion *ex vivo*.

Moreover, both *in vivo* and *ex vivo*, we show that prolonged hypoxia decreases the engagement of myogenic cells towards differentiation, as previously shown by others^18,20,47^. Stinkingly, the differentiation defect upon sustained hypoxia is fully restored upon progressive reoxygenation *ex vivo*, underlying the importance of transient hypoxia and reoxygenation in properly orchestrating the myogenic program *in vivo*. We also establish that prolonged hypoxia (1%O_2_) directly alters the fusion capacity of MuSCs with the formation of smaller myotubes *in vitro*, in agreement with previous studies^25,45,46^. While the dynamic of hypoxia during tissue repair has been overlooked, here we demonstrate that transient hypoxia followed by progressive reoxygenation rescues the fusion capacity of myogenic cells. These data suggest that blocking reoxygenation upon regeneration leads to a hypotrophic phenotype due to an impairment of both myogenic differentiation and fusion abilities.

We also establish, on single myofiber experiments, that prolonged hypoxia (1%O_2_) promotes MuSC self-renewal, as previously reported^16,48^. Moreover, transient hypoxia followed by progressive reoxygenation also favor MuSC self-renewal but to a lesser extent, underlying the crucial role of transient hypoxia for the maintenance of stemness in MuSCs.

At the molecular level, we confirm that the return into quiescence of MuSCs relies on HIF-1α stabilization, since it is lost in mice with MuSC-specific invalidation of HIF-1α, as previously demonstrated^15,49^. Conversely, the role of HIF signaling in modulating MuSC differentiation is still debated in literature since many *in vitro* studies, in line with ours, attest that hypoxia limits the commitment of MuSCs towards differentiation, independently of HIF-1α^20,22,23,25,50^. Accordingly, we demonstrate that under sustained hypoxia, MuSC-specific HIF-1α-null mice exhibit the same hypotrophic regenerating myofiber phenotype than CTRL mice, indicating that prolonged hypoxia delays skeletal muscle repair independently of HIF signalling. This result is in sharp contrast with the two main studies demonstrating the essential role of HIF-1α during muscle repair^21,49^. One explanation for this divergent result lies in the use, in their studies of two different Pax7-CreERT2 mouse lines to conditionally invalidate HIF-1α, that have both been shown to have technical limitations^51^. In our study, we used the Pax7-CreERT2 mouse line from the Kardon lab, which is the most used model to induce efficient recombination of floxed alleles in PAX7+ MuSCs^51^.

To decipher the mechanisms independent of HIF by which prolonged hypoxia slows MuSC differentiation and fusion during muscle repair, we performed a bulk RNAseq on PAX7^+^ cells freshly isolated from CTX-induced TAs of mice exposed to normoxia or hypoxia, at 7 dpi. Surprisingly, one of the most enriched pathways in our transcriptomic analysis was the circadian rhythm pathway. We identified a downregulation of the circadian clock activators and a parallel upregulation of the circadian clock inhibitor in PAX7+ cells exposed to prolonged hypoxia. This result echoes a recent study showing that exposure to microgravity, another atmospheric environmental factor is able to modulate MuSC fate and muscle physiology by altering the clock network, independently of light/dark cycle^52^. In the muscle field, multiple actors of the clock machinery have been shown to regulate MuSC fate. Among these actors, the transcription factor Bmal1 (Brain and Muscle ARNT-Like 1) has been shown to regulate MYOD expression and myofiber formation to support skeletal muscle homeostasis and repair^53,54^. In the same line, downstream effectors of circadian rhythm, such as PER1 and PER2 seems essential for myoblast differentiation and muscle regeneration^55^, whether the role of CRY2 remains debated^56,57^. Indeed, while Hao and colleagues demonstrated that whole-muscle *CRY2* deletion enhance muscle repair^56^, Lowe and colleague showed that *CRY2* loss in PAX7+ cells leads to smaller myotubes and impairs muscle regeneration as a consequence of inefficient fusion process^57^. Although muscle oxygen level was not addressed in the latter study, their results echo ours. Strikingly, in our transcriptomic analysis, the highest upregulated circadian clock gene in myogenic cells maintained under prolonged hypoxia is the circadian clock negative regulator, *Nr1d1* (Rev-ERBα). Its role on myogenesis in resting or regenerating muscles remains controversial. In healthy muscle, Rev-ERBα has been shown to promote *in vitro* myoblast differentiation^58^ and positively regulate myofiber size and muscle mass *in vivo*^59^. On the opposite, other studies suggest that Rev-ERBα inhibits myogenic differentiation *in vitro* by decreasing MyoD expression^60^ or by inactivating Wnt pathway^61^. Furthermore, inhibition of Rev-ERBα has been shown to ameliorate muscular dystrophy in mice, by stimulating myoblast differentiation and enhancing myofiber formation^62,63^. Since dystrophic muscles display marked microvasculature alterations, possibly leading to a hypoxic environment^64^, the impact of Rev-ERBα on myogenesis may be dependent on intramuscular oxygen level. In favor of this argument, our data show that maintaining the mice under prolonged hypoxia during muscle repair increases Rev-ERBα in myogenic cells and that pharmacological inhibition of Rev-ERBα restores the fusion capacity of MuSCs under prolonged hypoxia. Reciprocally, incubation of myogenic cells with a Rev-ERBa agonist reproduces *in vitro* the inhibitory impact of hypoxia on myogenic cell fusion. Our data is the first to demonstrate a causal link between low-oxygen environment, increased Rev-ERBα and altered myogenesis.

To conclude, our work highlights the critical role of transient hypoxia followed by progressive muscle reoxygenation for proper skeletal muscle repair, involving Rev-ERBα/circadian clock pathway.

## METHODS

### Mouse models

The mice used in this study were employed and maintained on C57BL/6N background: C57BL/6N (Janvier Labs^®^), *Tg:Pax7-nGFP* ^65^, *Pax7^CreERT^*^2^*^/+^* ^66^ and *Hif-1α^flox/flox^* ^67^. To induce recombinaison, five week-old *Pax7^CreERT^*^2^*^/+^;HIF-1α^flox/flox^* mice (HIF cKO) were fed with low-phytoestrogen tamoxifen diet (Altromin 1324P, Genetsil) for 10 consecutive days. Mice fed with normal diet have been used as control (HIF CTRL). Specific forward and reverse primers used for genotyping all the mice models are listed in *supplemental Table 1*.

Animals were handled according to national and European community guidelines and procedures were approved by the ethics committee at the French Ministry (Project No: 16-062). Mice were exposed to common 12h light/12h as day-night cycle. All samples were collected between 9 and 10 a.m.

### *In vivo* experimental design and muscle fixation

The *Tibialis Anterior* (TA) muscle was intramuscularly injected with 50μl of cobra venon called Cardiotoxin (CTX) (10μM, Latoxan laboratory; # L8102) on mice under isofluorane anesthesia. Then, mice have been placed in a normoxic environment at 21%O_2_ or in a normobaric hypoxic chamber (Biospherix) at 10%O_2_ inhaled to induce a systemic hypoxia during 5, 7, 10, 14 or 28 days post-CTX injury. All mice were exposed to 12h light/12h dark cycle and were subjected to *ad libitum* normal chow feeding and water. All the samples were collected in the morning.

In some experiments, mice have been intraperitoneally injected with a hypoxia probe named Pimonidazole (100μl; 60mg/kg; Hypoxyprobe-1^TM^; #HP1-100Kit) 30min before sacrifice. Pimonidazole, also known as 2-nitroimidazole, forms adducts with thiol groups of proteins, peptides and amino acids of cells under hypoxic stress (pO_2_ <10 mmHg) that can be detected by immunostaining.

At the end of the study, adult non-injured and injured TA muscles were harvested, immediately frozen in liquid-nitrogen-cooled isopentane and sectioned transversely at 8μm. Muscle sections were post-fixed with 4%PFA, 20 min at room temperature, washed with 1x PBS (3 times) before immunostaining as described in « Immunostaining on cells, sections and myofibers ».

### SUnSET assay

SUrface SEnsing of Translation assay was performed to detect changes in protein synthesis in whole TA muscles^28^ of mice placed in a normoxic (21%O_2_) or hypoxic (10%O_2_) during skeletal muscle repair. Mice were intraperitonealy injected with puromycin dihydrochloride (Sigma, P7255) at a dose of 40 nM/g of body weight, 30min before sacrifice. The non-injured and CTX-injured TA muscles were harvested and snap frozen in liquid nitrogen and stored at -80°C until further use described in “Protein extraction and Western-Blot”.

### Plasma hematocrit evaluation

Blood samples were collected through cardiac puncture from mice under isoflurane anesthesia in capillary tubes (Hirschmann^®^, #9100175). Tubes were centrifuged during 3 min at 10000rpm (Hawksley Micro-Haematocrit centrifuge) and the hematocrit levels were evaluated by measuring the proportion of red blood cells compared to total volume using an abacus.

### Isolation and culture of single myofibers

Single myofibers were isolated from EDL muscles following the previously described protocol^7^. Briefly, EDL muscles were dissected tendons to tendons and digested in a filtered solution of 0.2% collagenase (Sigma^®^; #C0130) in Dulbecco’s modified Eagle’s medium 4.5g/L glucose 1x DMEM-Glutamax^TM^-1 (Gibco^TM^; #31966), 1% Penicillin/Streptomycin (Gibco^TM^; #15140) for 1h30 at 37°C. After connective tissue digestion, mechanical dissociation was performed to release individual myofibers that were then transferred to serum-coated Petri dishes for 20 min. Single myofibers were either immediately fixed in 4% PFA for 10 min before immunostaining or maintained for 72h in myofiber growth medium containing 4.5g/L glucose 1x DMEM-Glutamax^TM^-1 (Gibco^TM^; #31966), 1% Penicillin/Streptomycin (Gibco^TM^; #15140), 20% fetal bovine serum (FBS; Gibco^TM^; #10270) and 1% chicken embryo extract (CEE; MP Biomedical; CE-650-J) at 37°C and 5%CO_2_. For 72h, myofibers were cultivated either at 8%O_2_ (physioxia), 1%O_2_ (hypoxia), or 1-8%O_2_ (transient hypoxia followed by progressive reoxygenation: 1%O_2_ for 18h followed by 2%O_2_ for 12h, 4%O_2_ for 12h, 6%O_2_ for 12h and finally 8%O_2_ for 18h). Myofibers were then recovered, and fixed with 4% PFA before immunostaining for proliferation, differentiation and quiescent markers, as described in « Immunostaining on cells, sections and myofibers ».

### Muscle enzymatic dissociation

Adult hindlimb and back muscles were dissected, minced and incubated with a mix of 2.4U/ml Dispase II (Roche, #04942078001) and 100μg/ml collagenase A (Roche, #11088793001) in 4.5g/L glucose 1x DMEM (Gibco^TM^; #11995) at 37 °C on a shaking plate for 2 h. The muscle suspension was filtrated through 100-μm and 70-μm cell strainers and then spun at 50g for 10min at 4°C to remove large tissue fragments. Then, the supernatant was collected, filtered through a 40-μm cell strainer and washed by centrifugation at 700g for 10 min at 4°C. The final pellet has been resuspended in cold 4.5g/L glucose 1xDMEM supplemented with 0.2% bovine serum albumin (BSA, Sigma-Aldrich). The muscle cell suspension was used either to sort MuSCs by fluorescence-activated cell sorting (FACS)-sorting or directly seeded on Matrigel coated dishes, as described in “MuSC isolation by flow cytometry”.

### MuSC isolation by flow cytometry

MuSCs were sorted from dissociated muscles with the cell sorter BD Influx Sorp (BD Biosciences) using either the GFP reporter for *Tg:Pax7-nGFP* mice or a cell surface labeling strategy for C57BL/6N, HIF cKO and HIF CTRL mice using anti-mouse CD45-PE-Cy7, anti-Ter119-PE-Cy7, anti-mouse CD34-BV421, anti-mouse Sca1-FITC (all from BD Biosciences, cf. *Supplemental Table 2*) and anti-mouse integrin-α7-A700 (R&D Systems, cf. *Supplemental Table 2*). Gating strategy for MuSC isolation using cell surface markers was as followed: Ter119^-^, CD45^-^, CD34^+^, Sca1^-^ and gating on the cell fraction, integrin-α7^+^.

### Muscle stem cell culture

For nuclear HIF-1α translocation experiments, freshly isolated MuSCs were cultured during 72h at 21%O_2_ and then stimulated for 3h at 20%O_2_ (normoxia), 1%O_2_ (hypoxia) or in presence with 1mM Dimethyloxalylglycine (DMOG; Thermo Fisher; #D3695), a pharmacological mimetic of hypoxia. Cells were fixed with 4% PFA before immunostaining for HIF-1α protein, as described in « Immunostaining on cells, sections and myofibers ».

For myoblastic fusion experiments, freshly isolated FACS-sorted MuSCs were plated on 8-well PCA on glass detachable plate (Sarsted, #94.6140.802) coated with Matrigel and cultured in proliferating medium containing 4,5g/L glucose 1x DMEM-Glutamax^TM^-1 (Gibco^TM^; #31966), 1% HEPES 1M (Gibco^TM^; #15630), 20% FBS (Gibco^TM^; #10270), 5% horse serum (Gibco^TM^; #16050) and primocin (100 µg/ml, Invitrogen) at 37°C and 5% CO_2_. Cells were maintained in culture for 72h at 8%O_2_ (physioxia), 1%O_2_ (prolonged hypoxia) or 1-8%O_2_ (transient hypoxia followed by progressive reoxygenation). MuSCs were seeded at 20 000 cells/cm^2^ for 8% and 1-8%O_2_ exposure and at 30 000 cells/cm^2^ for 1%O_2_ exposure. In some experiments, the Rev-Erbα antagonist SR8278 (5μM, MedChemExpress, #HY-14415), Rev-Erbα antagonist SR9009 (5μM, Sigma-Aldrisch, #554726) or theirs vehicles were added every 24h in the proliferation medium of MuSCs for 72h. After three days of culture, cells were fixed with 4% PFA and immunostained for cell fusion analysis, as described in « Immunostaining on cells, sections and myofibers ».

In some others experiments, freshly isolated MuSCs were plated at low density 10 000 cells/cm^2^ in 8-well PCA plate coated with Matrigel, cultured during 6h in proliferating growth medium describe above and culture for an additional 42h in a differentiation medium containing 4.5g/L glucose 1x DMEM-Glutamax^TM^-1 (Gibco^TM^; #31966), 1% Penicillin/Streptomycin (Gibco^TM^; #15140) and 5% horse serum (HS, Gibco^TM^; #16050) at 37°C, 5%CO_2_ at 8%O_2_, in order to favor the synchronous differentiation of myobastic cells without fusion. Differentiated cells were then trypsinized and plated at high-density (75 000 cells/cm^2^) in Matrigel coated 8-well plates, maintained for 6h in proliferation growth medium at 8%O_2_ and for 72h in differentiation medium either at 8%O_2_, 1%O_2_ or 1-8%O_2_ as described above. This protocol was adapted from B. Chazaud’s lab (REF). Cells were then fixed with 4% PFA and immunostained for cell fusion analysis, as described in « Immunostaining on cells, sections and myofibers ».

### Immunostaining on cells, sections and myofibers

Following PFA fixation, cells, muscles section or myofibers were washed three times with 1X PBS, then permeabilized and blocked at the same time with buffer containing 0.025% Tween20 for myofibers and 0.2% Triton X-100 for cells and sections. For PAX7 staining, sections were permeabilized with cold methanol, antigen retrieval was performed in boiling 10 mM citrate buffer pH 6 (Dako; #S2369) for 30 min and sections were incubated 30 min with Fab antibody. Samples were then incubated overnight at 4°C in 2.5% BSA in 0.025% Tween20 buffer with primary antibodies listed in *Supplemental table 2.* Samples were washed 3 times with 0.025% PBS-Tween20 and incubated with Alexa-conjugated secondary antibodies listed in *Supplemental Table 2* (Life Technologies, 1/1000e) for 1 h. After washing 3 times with 1X PBS, DAPI (Sigma, 1/5000e) was added for 5 min at room temperature. Samples were washed 3 times with 1X PBS and slides were mounted with Fluoromount-G medium (Interchim). Confocal images were acquired with a Zeiss LSM800 confocal (Zeiss) for representative pictures and analyzed with Zen Blue 2.0 software. Counting was performed using ImageJ (version 1.47 v; National Institutes of Health, USA, https://imagej.nih.gov/ij/) or under the Zeiss AxioImager D1 microscope (Zeiss).

### RNA extraction and RTqPCR

Total RNA was extracted from whole muscles using using PureLink™-RNA Mini Kit™ (Invitrogen™, 12183018A) or from GFP^+^ myogenic cells isolated from injured TA using RNAqueous™-Micro Kit RNA Isolation ™ (Invitrogen™, AM1931) and cDNA synthesis was performed using SuperScript™ IV VILO™ Master Mix with ezDNase™ Enzyme, according to manufacturer’s instructions. RNA quality was assessed by spectrophotometry (DeNovix DS-11 FX spectrometer). RTqPCR was performed using the Veriti® 96-well Fast Thermal Cycler (Applied Biosystems™) and realtime qPCR was performed with the StepOnePlus real-time PCR system (Applied Biosystems™) using SYBR™ Green detection tools (Applied Biosystems™). Expression of each gene was normalized to *TATA Box Protein (TBP)* gene expression. Results are reported as relative gene expression (2^-ΔCT^) to *TBP*. Specific forward and reverse primers used in this study are listed in *Supplemental Table 3*. Transcripts of Ang-2 were quantified using Taqman probe Angpt2: Mm00507897_ml (Applied Biosystems).

### Protein extraction and Western-blot

Frozen TA muscles were minced on ice and lysed in RIPA buffer (Tris 50mM, NaCl 150mM, EDTA 1mM, Na_3_VO_4_ 1mM, DOC 0,5%, Triton 1%, SDS 1%, and protease inhibitor cocktail 1X, all from Sigma) in Lysing Matrix D tubes (MP). The samples were mixed twice during 20s in Hybaïd Ribolyser, centrifuged at 13000rpm for 5min at 4°C. The supernatants were transferred into eppendorf tubes, centrifuged again at 13000rpm for 5min at 4°C and collected in fresh tubes. Protein concentrations were quantified using Pierce™ BCA Protein Assay Kit (Thermo Fisher, 23227).

Lysates containing 30μg of proteins were mixed with Bolt^TM^ LDS sample buffer 4X (Thermo Fisher B0007) containing Dithiothreitol UltraPure™ (Thermo Fisher 15508-013), boiled at 95°C for 5min and electrophoresed on Bolt^TM^ 4-12% Bis-Tris Plus gel (Thermo Fisher, NW04122BOX) at 200V for 20min. Proteins were then transferred onto iBlot^TM^ 2 PVDF membrane (Thermo Fisher, IB24002) at 20V for 7min using iBlot^TM^ 2 Gel Transfer Device (Thermo Fisher, IB 21001). Membrane was stained with red Ponceau-S (Merck, 1.15927) to control the quantity of proteins deposited for each sample and then blocked for 45min with 5% milk prepared in TBST (Tris-buffered saline supplemented with 2% Tween 20) at room temperature. Membrane was incubated overnight at 4°C with specific mouse monoclonal primary antibody anti-puromycin (Merck, MABE343, 1:1000) and after wash, for 1h at room temperature with HRP-conjugated anti-mouse secondary antibody (Santa Cruz, sc-2005, 1/10 000). Between each step, the blots were washed twice for 15min each, with TBST. Chemiluminescence was detected using SuperSignal^TM^ West Pico Plus reagent (Thermo Fisher, 34577) and the images were acquired using Azure Biosystems c600 imaging system and cSeries Capture software. The quantification of western-blot was performed using Image Lab™ Software (Bio-Rad).

### RNA sequencing

RNA sequencing was performed on FACS-sorted myogenic cells from dissociated TA muscles of *Tg:Pax7-nGFP* mice exposed to normoxic or hypoxic environment for 7 days after the induction of CTX injury. Libraries were constructed using an Illumina Stranded mRNA Prep (Illumina, USA) following the supplier’s recommendations. Library quality and quantity were assessed using the 2100 Bioanalyzer system (Agilent Technologies) and the Qbit dsDNA HS kit (ThermoFischer). Sequencing was performed using Illumina NextSeq 2000 platform using on the a P2 cadridge for a target of 50M reads single-ends 100 bases per sample according to the manufacturer’s instruction. The RNA-seq analysis was performed with Sequana 0.11.0 (https://github.com/sequana/sequana_rnaseq). Briefly, reads were trimmed from adapters using cutadapt 3.4 then mapped to the genome assembly GRCm38 from Ensembl using STAR 2.7.3a ^68^. FeatureCounts 2.0.1 ^69^ was used to produce the count matrix, assigning reads to features using corresponding annotation GRCm38_92 from Ensembl with strand-specificity information. Quality control statistics were summarized using MultiQC 1.10.1. Clustering of transcriptomic profiles were assessed using a Principal Component Analysis (PCA). Differential expression testing was conducted using DESeq2 library 1.24.0 ^70^. The normalization and dispersion estimation were performed with DESeq2 using the default parameters; statistical tests for differential expression were performed applying the independent filtering algorithm. A generalized linear model was set in order to test for the differential expression between conditions. Raw p-values were adjusted for multiple testing according to the Benjamini and Hochberg procedure and genes with an p-value adjusted lower than 0.05 were considered differentially expressed. Gene set enrichment analysis was performed using the fisher statistical test for the over-representation of up-regulated genes.

### Statistics

Results are presented as mean ± SEM. All statistical analysis and graphs were performed using GraphPad Prism^®^ Software (version 8.0). Student test (also called T-test) was used to compare two groups. For multiple comparisons, we used one-way ANOVA followed by Dunnet or Tukey’s post-test or two-way ANOVA followed by Sidak’s post-test. p<0.05 is considered as statistically significant.

## ACKNOWLEGMENTS

We would like to thank Aurélie Guguin, Adeline Henry and Odile Ruckebusch for their assistance with flow cytometry. We also thank Nathalie Chevalier, Nathalie Didier, Francesca Gattazzo and Béatrice Laurent for their help with the hypoxic incubators as well as Serge Adnot and Shariq Abid for the use of the normobaric hypoxic chamber. We thank Camille Laisne, Diana Gelperowic and Damien Fois for animal care and animal facility CDTA (TAAM, CNRS – UPS44). The RNAseq analysis was performed by E. Turc and E. Kornobis of Biomics Platform, C2RT, Institut Pasteur, Paris, France, supported by France Génomique (ANR-10-INBS-09) and IBISA. This work was supported by fundings from Association Française contre les Myopathies (AFM) via TRANSLAMUSCLE (PROJECT 19507) and MyoStemVasc (ANR-17-CE14-0018-01).

## AUTHORS CONTRIBUTION

Designed experiments: M.Q., M.G.; Performed experiments: M.Q., A.D.V., M.G., C.D., J.B., M.L., A.S., S.M. ; Generated mouse models: B.D.L and M.G. ; Wrote the manuscript: M.Q., A.D.V, M.G.; Funding Acquisition: M.G, F.R.; Supervision, M.G.

## REFERENCES

1. Pallafacchina G, Blaauw B, Schiaffino S. Role of satellite cells in muscle growth and maintenance of muscle mass. Nutr Metab Cardiovasc Dis. (2013).

2. Lepper C, Partridge TA, Fan CM. An absolute requirement for Pax7-positive satellite cells in acute injury-induced skeletal muscle regeneration. Development. 138(17):3639–46 (2011).

3. Seale P, Sabourin LA, Girgis-Gabardo A, Mansouri A, Gruss P, Rudnicki MA. Pax7 is required for the specification of myogenic satellite cells. Cell. 102(6):777–86 (2000).

4. Schmidt M, Schüler SC, Hüttner SS, von Eyss B, von Maltzahn J. Adult stem cells at work: regenerating skeletal muscle. Cell Mol Life Sci. 76(13):2559–2570 (2019).

5. Fukada, Si., Higashimoto, T. & Kaneshige, A. Differences in muscle satellite cell dynamics during muscle hypertrophy and regeneration. Skeletal Muscle 12, 17 (2022).

6. Leikina E, Gamage DG, Prasad V, Goykhberg J, Crowe M, Diao J, Kozlov MM, Chernomordik LV, Millay DP. Myomaker and Myomerger Work Independently to Control Distinct Steps of Membrane Remodeling during Myoblast Fusion. Dev Cell. 46(6):767–780 (2018).

7. Zammit PS, Golding JP, Nagata Y, Hudon V, Partridge TA, Beauchamp JR. Muscle satellite cells adopt divergent fates: a mechanism for self-renewal? J Cell Biol. 166(3):347–57 (2004).

8. Kostallari E, Baba-Amer Y, Alonso-Martin S, Ngoh P, Relaix F, Lafuste P, Gherardi RK. Pericytes in the myovascular niche promote post-natal myofiber growth and satellite cell quiescence. Development 142(7):1242–53 (2015).

9. Latroche C, Weiss-Gayet M, Muller L, Gitiaux C, Leblanc P, Liot S, Ben-Larbi S, Abou-Khalil R, Verger N, Bardot P, Magnan M, Chrétien F, Mounier R, Germain S, Chazaud B. Coupling between Myogenesis and Angiogenesis during Skeletal Muscle Regeneration Is Stimulated by Restorative Macrophages. Stem Cell Reports. 9(6):2018–2033 (2017).

10. Semenza GL. Oxygen sensing, hypoxia-inducible factors, and disease pathophysiology. Annu Rev Pathol. 9:47–71 (2014).

11. Lindholm ME, Rundqvist H. Skeletal muscle hypoxia-inducible factor-1 and exercise. Exp Physiol. 101(1):28–32 (2016).

12. Wagatsuma A, Kotake N, Yamada S. Spatial and temporal expression of hypoxia-inducible factor-1α during myogenesis in vivo and in vitro. Mol Cell Biochem. 347(1-2):145–55 (2011).

13. Rhoads RP, Johnson RM, Rathbone CR, Liu X, Temm-Grove C, Sheehan SM, Hoying JB, Allen RE. Satellite cell-mediated angiogenesis in vitro coincides with a functional hypoxia-inducible factor pathway. Am J Physiol Cell Physiol. 296(6):C1321–8 (2009).

14. Lemieux P, Birot O. Altitude, Exercise, and Skeletal Muscle Angio-Adaptive Responses to Hypoxia: A Complex Story. Front. Physiol. (2021).

15. Gustafsson MV, Zheng X, Pereira T, Gradin K, Jin S, Lundkvist J, Ruas JL, Poellinger L, Lendahl U, Bondesson M. Hypoxia requires notch signaling to maintain the undifferentiated cell state. Dev Cell. 9(5):617–28 (2005).

16. Liu W, Wen Y, Bi P, Lai X, Liu XS, Liu X, Kuang S. Hypoxia promotes satellite cell self-renewal and enhances the efficiency of myoblast transplantation. Development. 139(16):2857–65 (2012).

17. Verma M, Asakura Y, Murakonda BSR, Pengo T, Latroche C, Chazaud B, McLoon LK, Asakura A. Muscle Satellite Cell Cross-Talk with a Vascular Niche Maintains Quiescence via VEGF and Notch Signaling. Cell Stem Cell. 23(4):530–543 (2019).

18. Chaillou T, Lanner JT. Regulation of myogenesis and skeletal muscle regeneration: effects of oxygen levels on satellite cell activity. FASEB J. 30(12):3929–3941 (2016).

19. Cirillo F, Resmini G, Ghiroldi A, Piccoli M, Bergante S, Tettamanti G, Anastasia L. Activation of the hypoxia-inducible factor 1α promotes myogenesis through the noncanonical Wnt pathway, leading to hypertrophic myotubes. FASEB J. 31(5):2146–2156 (2017).

20. Zhang Z, Zhang L, Zhou Y, Li L, Zhao J, Qin W, Jin Z, Liu W. Increase in HDAC9 suppresses myoblast differentiation via epigenetic regulation of autophagy in hypoxia. Cell Death Dis. 10(8):552 (2019).

21. Majmundar AJ, Lee DS, Skuli N, Mesquita RC, Kim MN, Yodh AG, Nguyen-McCarty M, Li B, Simon MC. HIF modulation of Wnt signaling regulates skeletal myogenesis in vivo. Development. 142, 2405–2412 (2015).

22. Langley B, Thomas M, Bishop A, Sharma M, Gilmour S, Kambadur R. Myostatin inhibits myoblast differentiation by down-regulating MyoD expression. J Biol Chem. 277(51):49831–40 (2002).

23. Wang C, Liu W, Liu Z, Chen L, Liu X, Kuang S. Hypoxia Inhibits Myogenic Differentiation through p53 Protein-dependent Induction of Bhlhe40 Protein. J Biol Chem. 290(50):29707–16 (2015).

24. Di Carlo A, De Mori R, Martelli F, Pompilio G, Capogrossi MC, Germani A. Hypoxia inhibits myogenic differentiation through accelerated MyoD degradation. J. Biol. Chem. 279,16332–16338 (2004).

25. Majmundar AJ, Skuli N, Mesquita RC, Kim MN, Yodh AG, Nguyen-McCarty M, Simon MC. O(2) regulates skeletal muscle progenitor differentiation through phosphatidylinositol 3-kinase/AKT signaling. Mol. Cell. Biol. 32, 36–49 (2012).

26. Hardy D, Besnard A, Latil M, Jouvion G, Briand D, Thépenier C, Pascal Q, Guguin A, Gayraud-Morel B, Cavaillon JM, Tajbakhsh S, Rocheteau P, Chrétien F. Comparative Study of Injury Models for Studying Muscle Regeneration in Mice. PLoS One. 11(1):e0147198 (2016).

27. Varia MA, Calkins-Adams DP, Rinker LH, Kennedy AS, Novotny DB, Fowler WC Jr, Raleigh JA. Pimonidazole: a novel hypoxia marker for complementary study of tumor hypoxia and cell proliferation in cervical carcinoma. Gynecol Oncol. 71(2):270–7 (1998).

28. Goodman CA, Mabrey DM, Frey JW, Miu MH, Schmidt EK, Pierre P, Hornberger TA. Novel insights into the regulation of skeletal muscle protein synthesis as revealed by a new nonradioactive in vivo technique. FASEB J. 25(3):1028–39 (2011).

29. Relaix F, Machado L. Waking up muscle stem cells: PI3K signalling is ringing. EMBO J. 37(8):e99297 (2018).

30. Kok HJ, Barton ER. Actions and interactions of IGF-I and MMPs during muscle regeneration. Seminars in Cell & Developmental Biology 119: 11–22 (2021).

31. Pircher T, Wackerhage H, Aszodi A, Kammerlander C, Böcker W, Saller MM. Hypoxic Signaling in Skeletal Muscle Maintenance and Regeneration: A Systematic Review. Front Physiol. (2021).

32. Drouin G, Couture V, Lauzon MA, Balg F, Faucheux N, Grenier G. Muscle injury-induced hypoxia alters the proliferation and differentiation potentials of muscle resident stromal cells. Skelet Muscle. 9(1):18 (2019).

33. Xie L, Yin A, Nichenko AS, Beedle AM, Call JA, Yin H. Transient HIF2A inhibition promotes satellite cell proliferation and muscle regeneration. J Clin Invest. 128(6):2339–2355 (2018).

34. Verma M, Asakura Y, Murakonda BSR, Pengo T, Latroche C, Chazaud B, McLoon LK, Asakura A. Muscle Satellite Cell Cross-Talk with a Vascular Niche Maintains Quiescence via VEGF and Notch Signaling. Cell Stem Cell. 23(4):530–543.e9 (2018).

35. Befani C, Liakos P. The role of hypoxia-inducible factor-2 alpha in angiogenesis. J Cell Physiol. 233(12):9087–9098 (2018).

36. Deldicque L, Francaux M. Acute vs. chronic hypoxia: What are the consequences for skeletal muscle mass? Cellular and Molecular Exercise Physiology. 2 (2013).

37. Schiaffino S, Dyar KA, Ciciliot S, Blaauw B, Sandri M. Mechanisms regulating skeletal muscle growth and atrophy. FEBS J. 280(17):4294–314 (2013).

38. Gosker HR, Zeegers MP, Wouters EF, Schols AM. Muscle fibre type shifting in the vastus lateralis of patients with COPD is associated with disease severity: a systematic review and meta-analysis. Thorax. 62(11):944–9 (2007).

39. McGuigan MR, Bronks R, Newton RU, Sharman MJ, Graham JC, Cody DV, Kraemer WJ. Muscle fiber characteristics in patients with peripheral arterial disease. Med Sci Sports Exerc. 33(12):2016–21 (2001).

40. Shiragaki-Ogitani M, Kono K, Nara F, Aoyagi A. Neuromuscular stimulation ameliorates ischemia-induced walking impairment in the rat claudication model. J Physiol Sci. 69(6):885–893 (2019).

41. Liu F, Fry CS, Mula J, Jackson JR, Lee JD, Peterson CA, Yang L. Automated fiber-type-specific cross-sectional area assessment and myonuclei counting in skeletal muscle. J Appl Physiol (1985). 115(11):1714–24 (2013).

42. Endo Y, Zhu C, Giunta E, Guo C, Koh DJ, Sinha I. The Role of Hypoxia and Hypoxia Signaling in Skeletal Muscle Physiology. Adv Biol (Weinh). e2200300 (2023).

43. Ren H, Accili D, Duan C. Hypoxia converts the myogenic action of insulin-like growth factors into mitogenic action by differentially regulating multiple signaling pathways. Proc Natl Acad Sci USA. 107(13):5857–62 (2010).

44. Urbani L, Piccoli M, Franzin C, Pozzobon M, De Coppi P. Hypoxia increases mouse satellite cell clone proliferation maintaining both in vitro and in vivo heterogeneity and myogenic potential. PLoS One. 7, e49860 (2012).

45. Launay T, Hagström L, Lottin-Divoux S, Marchant D, Quidu P, Favret F, Duvallet A, Darribère T, Richalet JP, Beaudry M. Blunting effect of hypoxia on the proliferation and differentiation of human primary and rat L6 myoblasts is not counteracted by Epo. Cell Prolif. 43, 1–8 (2010).

46. Hidalgo M, Marchant D, Quidu P, Youcef-Ali K, Richalet JP, Beaudry M, Besse S, Launay T. Oxygen modulates the glutathione peroxidase activity during the L6 myoblast early differentiation process. Cell. Physiol. Biochem. 33, 67–77 (2014).

47. Bentzinger F, Wang YX, Dumont NA, Rudnicki MA. Cellular dynamics in the muscle satellite cell niche. EMBO Rep. 14(12):1062–72 (2013).

48. Wagatsuma A, Arakawa M, Matsumoto H, Matsuda R, Hoshino T, Mabuchi K. Cobalt chloride, a chemical hypoxia-mimicking agent, suppresses myoblast differentiation by downregulating myogenin expression. Mol Cell Biochem. 470, 199–214 (2020).

49. Yang X, Yang S, Wang C, Kuang S. The hypoxia-inducible factors HIF1α and HIF2α are dispensable for embryonic muscle development but essential for postnatal muscle regeneration. J Biol Chem. 292(14):5981–5991 (2017).

50. Yun Z, Lin Q, Giaccia AJ. Adaptive myogenesis under hypoxia. Mol Cell Biol. 25(8):3040–55 (2005).

51. Mademtzoglou D, Geara P, Mourikis P, Relaix F. Pax7 haploinsufficiency impairs muscle stem cell function in Cre-recombinase mice and underscores the importance of appropriate controls. Stem Cell Res Ther. 14(1):294 (2023).

52. Malhan D, Yalçin M, Schoenrock B, Blottner D, Relógio A. Skeletal muscle gene expression dysregulation in long-term spaceflights and aging is clock-dependent. NPJ Microgravity. 9(1):30 (2023).

53. Andrews JL, Zhang X, McCarthy JJ, McDearmon EL, Hornberger TA, Russell B, Campbell KS, Arbogast S, Reid MB, Walker JR, Hogenesch JB, Takahashi JS, Esser KA. CLOCK and BMAL1 regulate MyoD and are necessary for maintenance of skeletal muscle phenotype and function. Proc Natl Acad Sci USA. 107(44):19090–5 (2010).

54. Chatterjee S, Yin H, Nam D, Li Y, Ma K. Brain and muscle Arnt-like 1 promotes skeletal muscle regeneration through satellite cell expansion. Exp Cell Res. 331(1):200–210 (2015).

55. Katoku-Kikyo N, Paatela E, Houtz DL, Lee B, Munson D, Wang X, Hussein M, Bhatia J, Lim S, Yuan C, Asakura Y, Asakura A, Kikyo N. Per1/Per2-Igf2 axis-mediated circadian regulation of myogenic differentiation. J Cell Biol. 220(7):e202101057 (2021).

56. Hao Y, Xue T, Liu SB, Geng S, Shi X, Qian P, He W, Zheng J, Li Y, Lou J, Shi T, Wang G, Wang X, Wang Y, Li Y, Song YH. Loss of CRY2 promotes regenerative myogenesis by enhancing PAX7 expression and satellite cell proliferation. MedComm. 4(1):e202 (2020).

57. Lowe M, Lage J, Paatela E, Munson D, Hostager R, Yuan C, Katoku-Kikyo N, Ruiz-Estevez M, Asakura Y, Staats J, Qahar M, Lohman M, Asakura A, Kikyo N. Cry2 Is Critical for Circadian Regulation of Myogenic Differentiation by Bclaf1-Mediated mRNA Stabilization of Cyclin D1 and Tmem176b. Cell Rep. 22(8):2118–2132 (2018).

58. Dadon-Freiberg M, Chapnik N, Froy O. REV-ERBα activates the mTOR signalling pathway and promotes myotubes differentiation. Biol Cell. 112(8):213–221 (2020).

59. Mayeuf-Louchart A, Thorel Q, Delhaye S, Beauchamp J, Duhem C, Danckaert A, Lancel S, Pourcet B, Woldt E, Boulinguiez A, Ferri L, Zecchin M, Staels B, Sebti Y, Duez H. Rev-erb-α regulates atrophy-related genes to control skeletal muscle mass. Sci Rep. 7(1):14383 (2017).

60. Downes, M., Carozzi, A. J. & Muscat, G. E. Constitutive expression of the orphan receptor, Rev-erbA alpha, inhibits muscle differentiation and abrogates the expression of the myoD gene family. Molecular endocrinology 9, 1666–1678 (1995).

61. Chatterjee S, Yin H, Li W, Lee J, Yechoor VK, Ma K. The Nuclear Receptor and Clock Repressor Rev-erbα Suppresses Myogenesis. Sci Rep. 9(1):4585 (2019).

62. Welch RD, Billon C, Kameric A, Burris TP, Flaveny CA. Rev-erbα heterozygosity produces a dose-dependent phenotypic advantage in mice. PLoS One. 15(5):e0227720 (2020).

63. Xiong X, Gao H, Lin Y, Yechoor V, Ma K. Inhibition of Rev-erbα ameliorates muscular dystrophy. Exp Cell Res. 406(2):112766 (2021).

64. Latroche C, Matot B, Martins-Bach A, Briand D, Chazaud B, Wary C, Carlier PG, Chrétien F, Jouvion G. Structural and Functional Alterations of Skeletal Muscle Microvasculature in Dystrophin-Deficient mdx Mice. Am J Pathol. 185(9):2482–94 (2015).

65. Sambasivan R, Gayraud-Morel B, Dumas G, Cimper C, Paisant S, Kelly RG, Tajbakhsh S. Distinct regulatory cascades govern extraocular and pharyngeal arch muscle progenitor cell fates. Dev Cell. 16(6):810–21 (2009).

66. Murphy MM, Lawson JA, Mathew SJ, Hutcheson DA, Kardon G. Satellite cells, connective tissue fibroblasts and their interactions are crucial for muscle regeneration. Development. 138(17):3625–37 (2011).

67. Ryan HE, Poloni M, McNulty W, Elson D, Gassmann M, Arbeit JM, Johnson RS. Hypoxia-inducible factor-1alpha is a positive factor in solid tumor growth. Cancer Res 60(15):4010–5 (2000).

68. Dobin A. Davis C.A. Schlesinger F. Drenkow J. Zaleski C. Jha S. Batut P. Chaisson M. Gingeras T.R. STAR: ultrafast universal RNA-seq aligner. Bioinformatics. 29: 15–21 (2013).

69. Liao Y. Smyth G.K. Shi W. featureCounts: an efficient general purpose program for assigning sequence reads to genomic features. Bioinformatics. 30: 923–930 (2014).

70. Love M.I. Huber W. Anders S. Moderated estimation of fold change and dispersion for RNA-seq data with DESeq2. Genome Biol. 15: 550 (2014).

