## Supplemental Figures and Tables for "Transient hypoxia followed by progressive reoxygenation is required for efficient skeletal muscle repair through Rev-ERBα modulation"

**SUPPLEMENTAL INFORMATION**

Supplemental information contains three figures and three tables.

**Supplemental figure 1.**

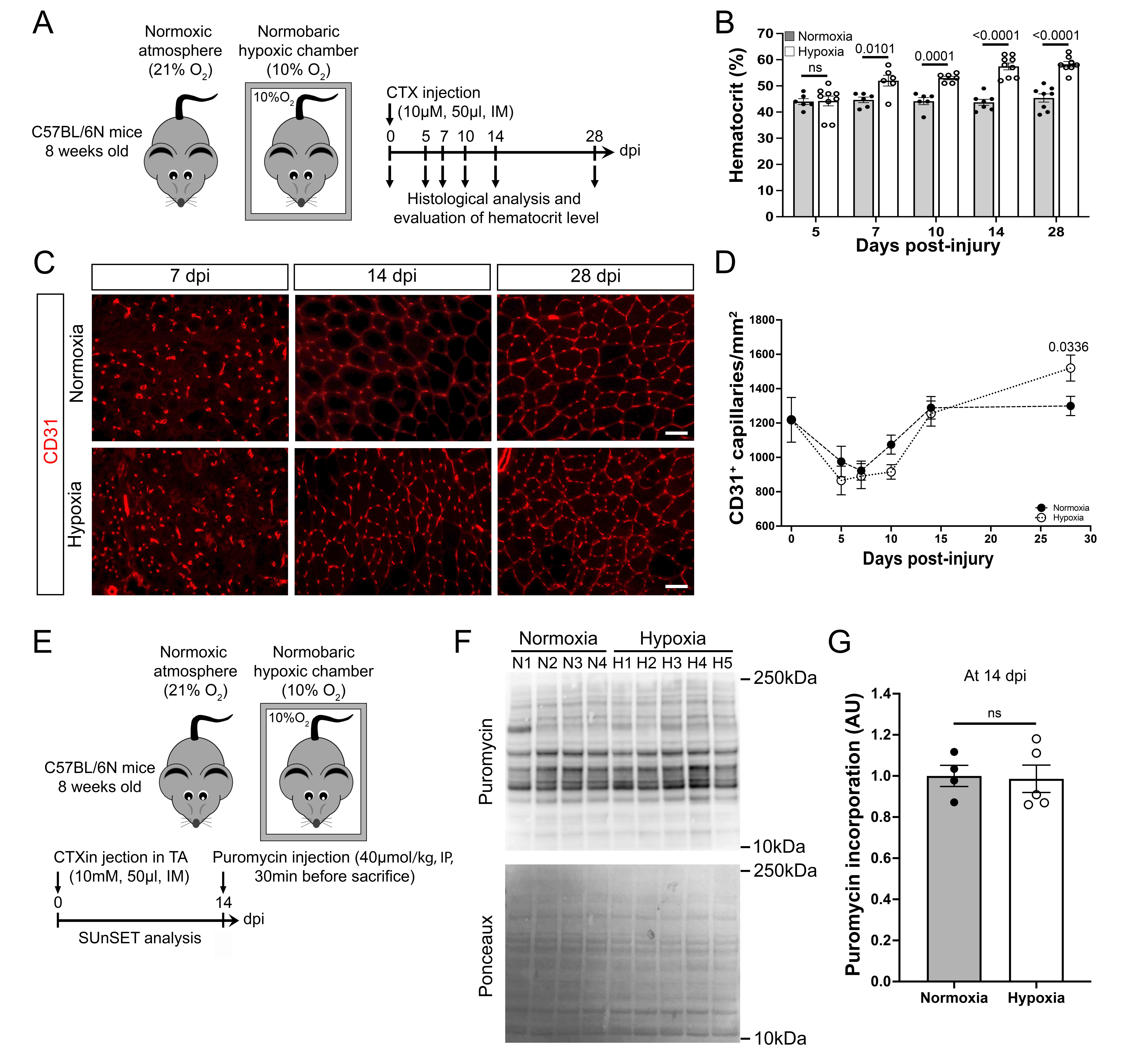

**Supplemental figure 1: Skeletal muscle revascularization following injury in the *Tibialis Anterior* muscle of adult mice under systemic hypoxia.**

**A.** Experimental design. Acute muscle injury was induced by cardiotoxin (CTX) intramuscular (IM) injection in the tibialis anterior (TA) of 8-week-old wild-type mice. Mice were then housed in standard normoxic atmosphere (21%O_2_) or in a normobaric hypoxic chamber (10%O_2_), from 0 to 28 days post-injury (dpi).

**B.** Evaluation of hematocrit (%) in mice exposed to normoxia or prolonged hypoxia at 5, 7, 14 and 28 days post-injury (dpi).

**C.** Representative pictures of CD31+ capillaries staining on cross-sections of CTX-injured TAs from mice placed under normoxia or prolonged hypoxia, from 0 to 28 dpi. Scale bar: 50μm.

**D.** Quantification of CD31^+^ capillaries/mm^2^ on cross-sections of CTX-injured TAs from mice placed under normoxia or prolonged hypoxia, from 0 to 28 dpi.

**E.** Experimental design. Acute muscle injury was induced by cardiotoxin (CTX) intramuscular (IM) injection in the tibialis anterior (TA) of 8-week-old wild-type mice. Mice were then housed in standard normoxic atmosphere (21%O_2_) or in a normobaric hypoxic chamber (10%O_2_), from 0 to 28 days post-injury (dpi). Intraperitoneal (IP) injection of puromycin were performed at 14dpi, 30min before mice sacrifice.

**F.** Representative images of membranes stained with puromycin and Ponceau from SUnSET analysis performed on protein extracts from whole CTX-injured TAs of mice housed in normoxia or prolonged hypoxia, at 14 dpi.

**G.** Quantitation of puromycin incorporation (AU) based on western-blot band intensity from 2.F.

**Statistics:** Results are expressed as means ± SEM.

For **B, D and G.** Unpaired t-test. n=6-9 per time point per group (B, D) and n=4 per atmosphere exposure (G).

**Supplemental figure 2.**

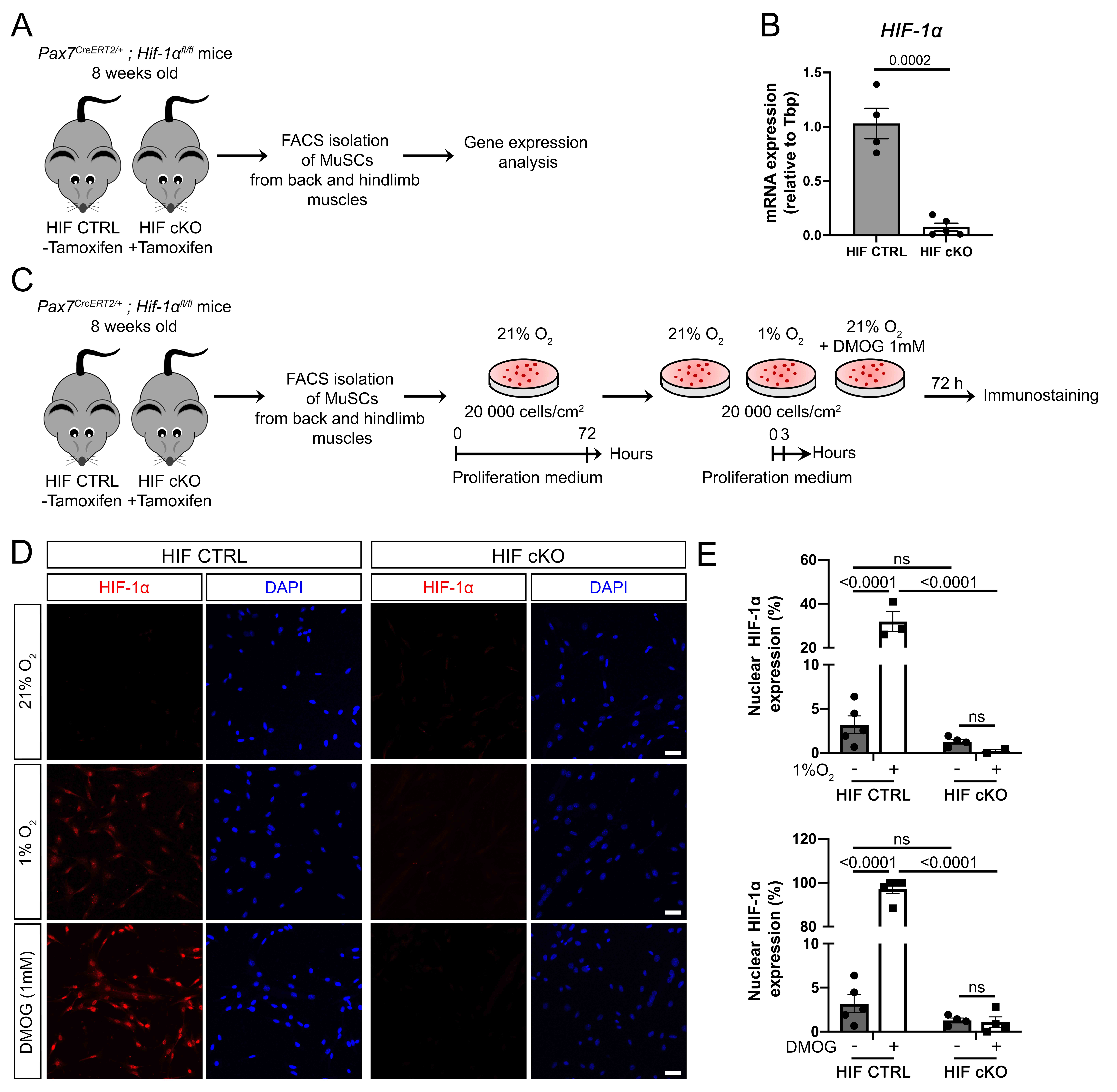

**Supplemental figure 2: Validation of HIF-1α knock-out in muscle stem cell mouse model**

**A.** Experimental design. Muscle stem cells (MuSCs) were isolated by FACS from HIF CTRL and HIF cKO mice. After lysis and mRNA extraction of isolated MuSCs, HIF-1α gene expression was analysed by quantitative Polymerase Chain Reaction (qPCR).

**B.** Histogram showing the relative expression *HIF-1α* normalized to *TBP* (Tata Box Protein) on FACS-sorted myogenic cell from muscles of HIF CTRL and HIF cKO mice.

This experiment validates the efficiency of HIF-1α deletion following tamoxifen-induced Cre ablation of HIF-1α in HIF cKO MuSCs compared to HIF CTRL in MuSCs.

**C.** Experimental design. MuSCs were isolated by FACS from back and hindlimb muscles of HIF CTRL and HIF cKO mice. Isolated MuSCs were cultured *in vitro* for 72h in normoxia (21%O_2_) and then placed for 3h in hypoxia (1%O_2_) or in normoxia (21%O_2_) with or without a pharmacological molecule allowing to mimic hypoxia DMOG (21%O_2_ + DMOG 1M).

**D.** Representative pictures of HIF-1α (red) and nuclei (DAPI, blue) staining on myogenic cells from HIF CTRL and HIF cKO mice, after 72h of culture under 21% O_2_ and 3h under 21%O_2_ with/without DMOG or 1%O_2_. Scale bar: 50μm.

**E.** Quantification of the nuclear translocation of HIF-1α (number of DAPI^+^ HIF-1α^+^ nuclei/total number of DAPI^+^ nuclei, %) in cultured myogenic cells from HIF CTRL or HIF cKO mice, after 72h of culture under 21% O_2_ and 3h under 21%O_2_ with/without DMOG or 1%O_2._ As expected, HIF-1α accumulates after hypoxic stimuli (i.e. hypoxia at 1%O_2_ and normoxic exposure (21%O_2_) with DMOG) into the nucleus of HIF CTRL myogenic cells while this nuclear translocation was abrogated in HIF-1α cKO myogenic cells.

**Statistics:** Results are expressed as means ± SEM.

For **B.** Unpaired t-test. n=4 per group.

For **E.** One-way ANOVA followed by Tukey’s post-test. n=2-5 per mice group per condition.

**Supplemental figure 3.**

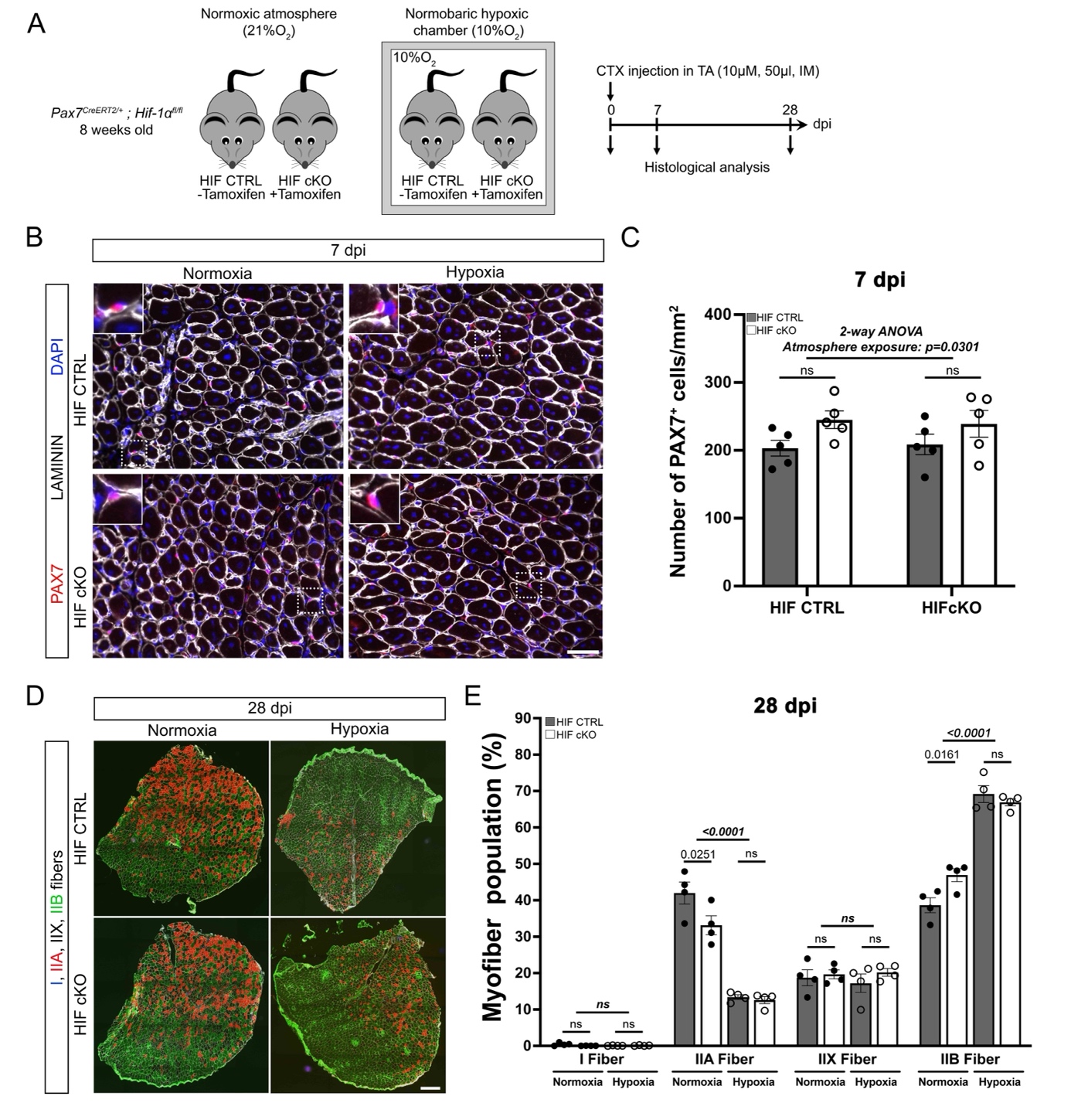

**Supplemental figure 3: Impact of conditional deletion of HIF-1α in muscle stem cells on muscle repair and MuSCs self-renewal.**

**A.** Experimental design. Experimental design. Acute muscle injury was induced by cardiotoxin (CTX) intramuscular (IM) injection in the tibialis anterior (TA) muscles from 8-week-old HIF CTRL and HIF cKO mice. Mice were then housed in standard normoxic atmosphere (21%O_2_) or in a normobaric hypoxic chamber (10%O_2_) from 0 to 28 days post-injury (dpi).

**B.** Representative pictures of PAX7 (red), LAMININ (white) and nuclei (DAPI, blue) staining on cross-sections CTX-injured TAs from HIF CTRL and HIF cKO mice exposed to normoxia or prolonged hypoxia, at 7 dpi. Scale bars: 50 μm (overviews) and 12.5 μm (insets).

**C.** Quantification of PAX7^+^ cells in CTX-injured TAs of HIF CTRL and HIF cKO mice exposed to normoxia or prolonged hypoxia, at 7 dpi.

**D.** Representative pictures of type-I myofibers (blue), type-IIA myofibers (red), type-IIB (green) and LAMININ (white) staining on cross-sections CTX-injured TAs of HIF CTRL and HIF cKO mice, at 28 dpi. Scale bar: 200 μm.

**E.** Quantification of the percentage of each myofiber type population of CTX-injured TAs injury of HIF CTRL and HIF cKO mice exposed to normoxia or prolonged hypoxia, at 28 dpi.

**Statistics:** Results are expressed as means ± SEM.

For **C and E.** 2-way ANOVA followed by Sidàk’s post-test. n=5 per mice group and per atmosphere exposure (C) and n=4 per mice per group and per atmosphere exposure (E).

**Supplemental Table 1.** List of specific forward and reverse primers used for genotyping.

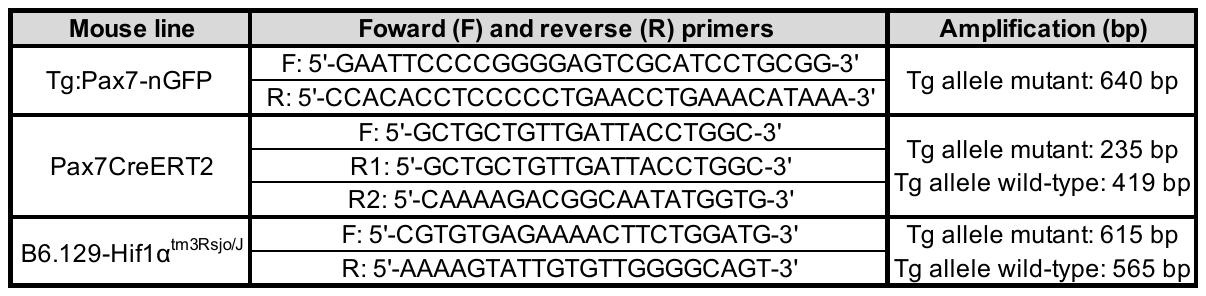

**Supplemental Table 2.** List of primary, secondary and coupled antibodies used for immunostaining, western-blot and FACS-sorting.

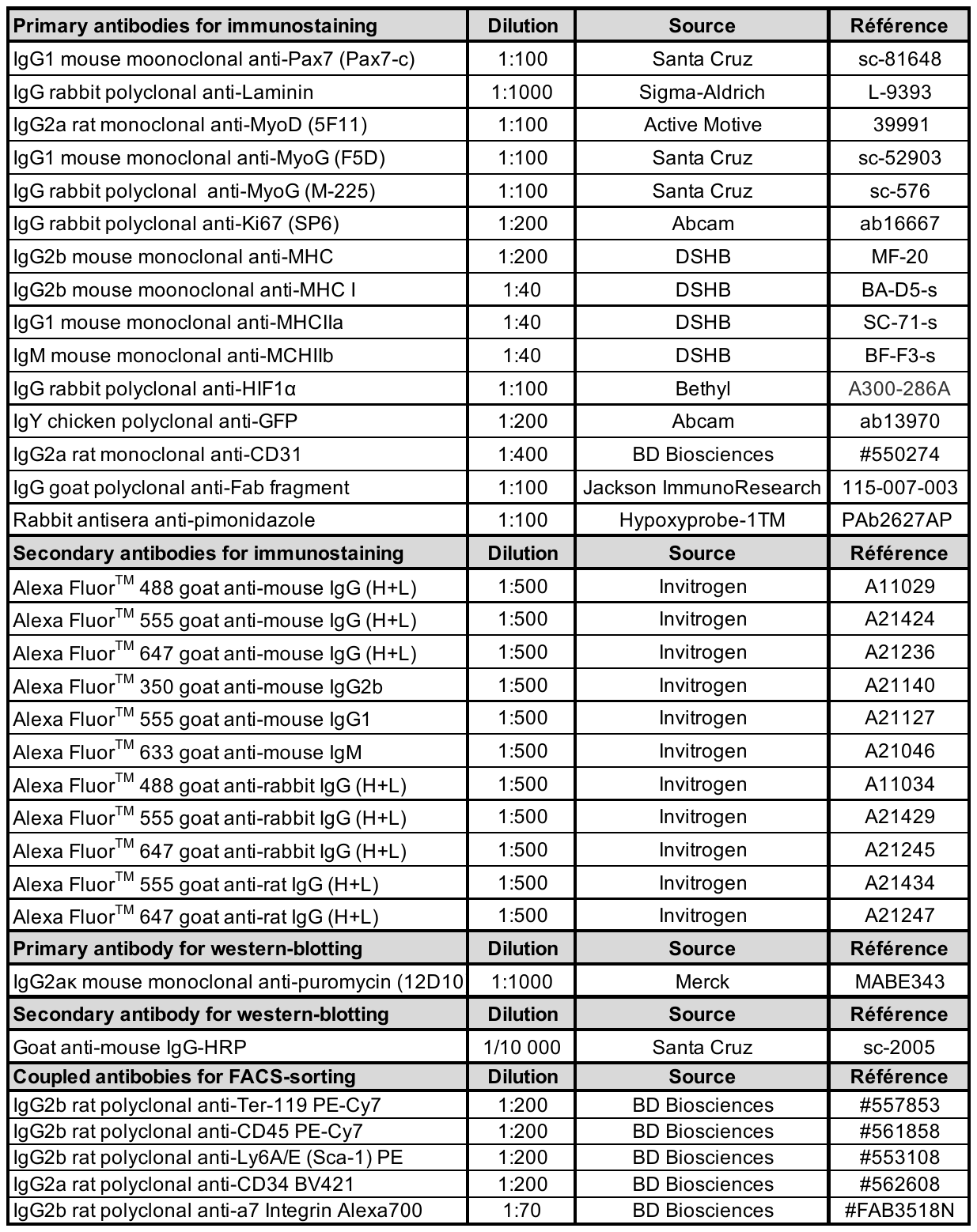

**Supplemental Table 3.** Specific forward and reverse primers used for RTqPCR

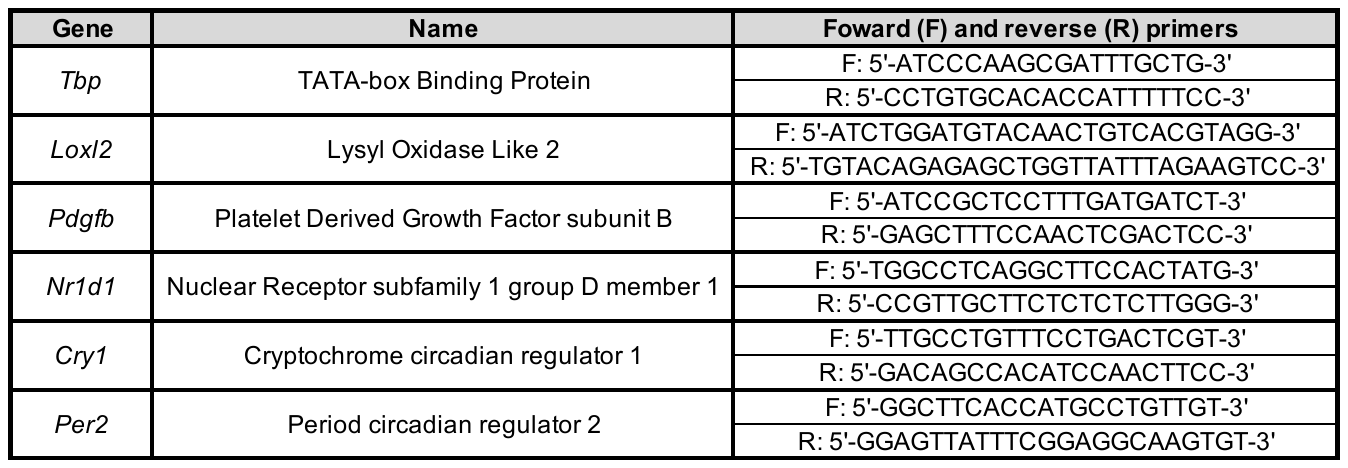
